# Unprocessed Genomic Uracil as a Source of DNA Replication Stress in Cancer Cells

**DOI:** 10.1101/2024.02.05.578390

**Authors:** Sneha Saxena, Parasvi S. Patel, Christopher S. Nabel, Ajinkya S. Kawale, Caroline R. Crosby, Matthew G. Vander Heiden, Aaron N. Hata, Lee Zou

## Abstract

Alterations of bases in DNA constitute a major source of genomic instability. It is believed that base alterations trigger base excision repair (BER), generating DNA repair intermediates interfering with DNA replication. Here, we show that genomic uracil, a common base alteration, induces DNA replication stress (RS) without being processed by BER. In the absence of uracil DNA glycosylase (UNG), genomic uracil accumulates to high levels, DNA replication forks slow down, and PrimPol-mediated repriming is enhanced, generating single-stranded gaps in nascent DNA. ATR inhibition in UNG-deficient cells blocks repair of uracil-induced gaps, increasing replication fork collapse and cell death. Notably, a subset of cancer cells harboring high levels of genomic uracil upregulate UNG2 to limit RS, and these cancer cells are hypersensitive to co-treatment with ATR inhibitors and drugs increasing genomic uracil. These results reveal unprocessed genomic uracil as an unexpected source of RS and a targetable vulnerability of cancer cells.

## Introduction

Complete and accurate DNA replication is essential for the maintenance of genomic integrity. However, the progression of DNA replication forks is often impeded by various types of barriers or interferences, resulting in DNA replication stress (RS) (Saxena and Zou, 2022; Zeman and Cimprich, 2014). Increased RS is commonly observed in cancer cells, which contributes to the genomic instability in cancer cells but also presents a vulnerability of cancer cells that can be targeted therapeutically (da Costa et al., 2022; Hopkins et al., 2022; Kotsantis et al., 2018; Macheret and Halazonetis, 2015). Understanding the sources of RS in cancer cells is critical for delineating the process of tumorigenesis and developing strategies to exploit RS in cancer therapy.

The common impediments or interferences to DNA replication forks can be divided into at least three classes. The first class comprises various types of physical barriers to replication forks, including bulky DNA adducts, DNA crosslinks, protein-DNA crosslinks, R-loops, transcription-replication conflicts, and others (Brickner et al., 2022; Cortez, 2019; Petermann et al., 2022). The second class includes various causes of insufficient dNTP supply, which limit the activity of DNA polymerases (Pai and Kearsey, 2017). The third class includes base alternations in DNA, such as those caused by guanine oxidation, cytosine deamination, TET-mediated cytosine demethylation, and misincorporation of uracil into DNA (Bauer et al., 2015). These base alterations trigger base excision repair (BER), generating abasic (AP) sites and DNA single-strand breaks (SSBs) as repair intermediates (Caldecott, 2020) (Fig. 1A). AP sites and SSBs, if not removed timely and properly, can interfere with replication forks and generate DNA double-stranded breaks (DSBs) (Fugger et al., 2021; Thompson and Cortez, 2020). Although base alterations are implicated in the generation of RS, it is still unclear whether this type of RS is always dependent on BER.

**Figure 1.**
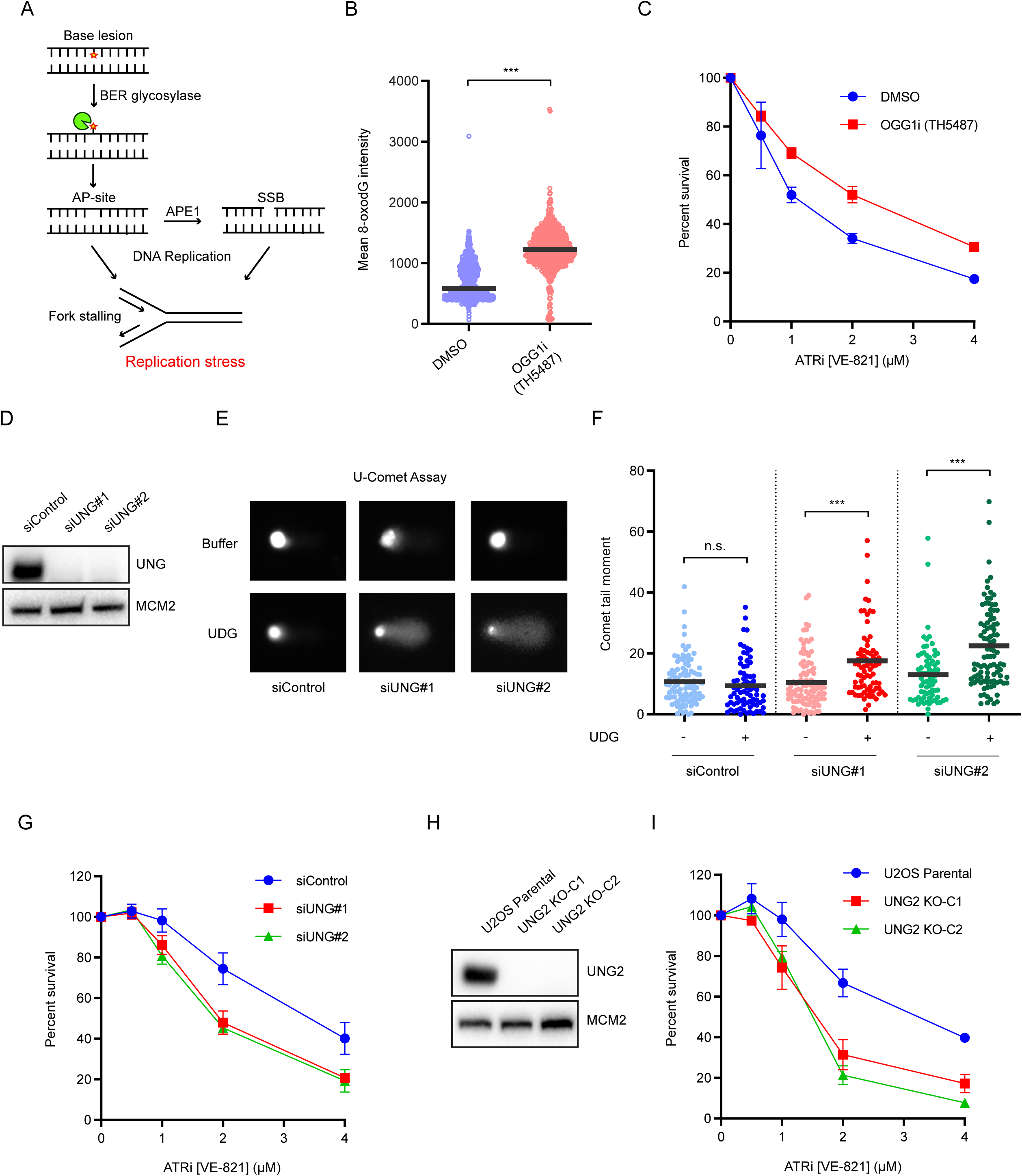
UNG2-mediated processing of genomic uracil reduces ATRi sensitivity. (A) Schematic representation of induction of replication stress by base excision repair (BER) intermediates—AP-sites and single strand DNA breaks (SSBs). (B) Quantification of 8-oxoG immunofluorescence in cells treated with DMSO or 10 µM TH5487 for 6 h (n>900 cells per condition). (C) U2OS cells were treated with DMSO or 10 µM TH5487 and indicated concentrations of ATRi (VE-821) for 5-7 days. Cell viability was determined using CellTiter-Glo and normalized to the cells untreated with ATRi under each condition for each condition. Data are shown as mean ± s.d. (n=2 independent experiments). (D) U2OS cells were transfected with the indicated siRNAs for 48 h. Levels of UNG in whole-cell extracts were analyzed by western blot. MCM2 is shown as the loading control. (E and F) Representative images for U-comet assay in U2OS cells transfected with indicated siRNAs. Dot plots represent comet tail moment in dividual cells, and bars display the mean in cell populations (n>75 cells per condition). One of two independent experiments is shown. (G) U2OS cells were transfected with indicated siRNAs and treated with indicated concentrations of ATRi (VE-821) for 5-7 days. Cell viability was determined using CellTiter-Glo and normalized to the cells untreated with ATRi under each condition. Data are shown as mean ± s.d. (n=2 independent experiments). (H) Levels of UNG2 in whole-cell extracts from wild-type U2OS cells and two independent *UNG2*^-/-^ knockout (KO) clones were analyzed by western blot. MCM2 is shown as the loading control. (I) WT U2OS or *UNG2* KO cells were treated with indicated concentrations of ATRi (VE-821) for 5-7 days. Cell viability was determined using CellTiter-Glo and normalized to the cells untreated with ATRi under each condition. Data are shown as mean ± s.d. (n=2 independent experiments).

In cancer cells, the levels of base alterations in DNA are often elevated by the increase of reactive oxygen species (ROS) (Szatrowski and Nathan, 1991), aberrant expression of AID and APOBEC cytidine deaminases (Petljak et al., 2022; Pettersen et al., 2015), and changes in dUTP and dTTP biogenesis (Berger et al., 2008). Thus, increased base alterations in DNA constitute an important source of RS in cancer cells and potentially provide an opportunity for targeted therapy. The ataxia telangiectasia-mutated and Rad3-related (ATR) kinase is a master regulator of the cellular response to RS, and it is critical for cancer cells to cope with RS and genomic instability (Hopkins et al., 2022; Saldivar et al., 2017; Zou and Elledge, 2003). Recently, ATR inhibitors (ATRi) have been successfully used to eliminate cancer cells under specific oncogene-induced RS (Konstantinopoulos et al., 2021; Thomas et al., 2021). However, it remains unclear whether ATRi can effectively kill cancer cells harboring high levels of base alterations and whether BER intermediates are important for ATRi sensitivity. Furthermore, whether different types of base alterations generate RS and confer ATRi sensitivity through the same or distinct mechanisms remains unknown.

## Results

### Processing of genomic uracil by UNG2 reduces ATRi sensitivity

To test whether base alterations induce replication stress and confer ATRi sensitivity through BER intermediates, we sought to ablate various DNA glycosylases, the enzymes that recognize different types of base alterations and initiate BER (Fig. 1A). We first inhibited OGG1, the DNA glycosylase that recognizes and processes 8-oxoG (oxidized guanine), in U2OS cells with the OGG1 inhibitor (OGG1i) TH5487 (Visnes et al., 2018). As expected, OGG1i increased the levels of 8-oxoG in cells (Fig. 1B), confirming that the conversion of 8-oxoG to AP site by OGG1, the first step of BER, was inhibited. OGG1i reduced the sensitivity of cells to two distinct ATRis, VE-821 and AZD6738 (Fig. 1C, S1A), supporting the idea that 8-oxoG confers ATRi sensitivity through BER.

Next, we tested whether the deoxyuracil (dU) in DNA also confers ATRi sensitivity through BER. Uracil in DNA is a common base alteration resulting from the misincorporation of dUTP during DNA replication or deamination of cytosine. The uracil glycosylases UNG1 and UNG2, which are encoded by two isoforms of the *UNG* gene, are the primary enzymes that recognize and process the uracil in DNA (Krokan et al., 2002). Whereas UNG1 localizes to mitochondria, UNG2 functions in the nucleus. We used two independent siRNAs, each targeting both isoforms of *UNG*, to deplete UNG in U2OS cells (Fig. 1D, S1B). To confirm the effects of UNG loss, we used a modified comet assay (U-comet) to measure uracil levels in genomic DNA (Fig. 1E). In this assay, cells were treated with recombinant uracil DNA glycosylase (UDG) before analyzed by the alkaline comet protocol, allowing detection of DNA breakage at sites of genomic uracil. As expected, UNG knockdown increased genomic uracil (Fig. 1E-F), which was associated with a concomitant reduction in AP sites (Fig. S1C), showing that the uracil in DNA was not efficiently processed in the absence of UNG. Surprisingly, however, knockdown of UNG increased the ATRi sensitivity of cells (Fig. 1G, S1D), suggesting that BER does not enhance ATRi sensitivity in this context. Expression of HA-tagged UNG2 significantly reversed the increase of ATRi sensitivity in UNG knockdown cells (Fig. S1E-F), confirming that the effect of UNG knockdown on ATRi sensitivity is largely, if not completely, attributed to UNG2 loss. To confirm the results from UNG knockdown cells and test whether UNG2 loss is responsible for increased ATRi sensitivity, we generated 2 independent UNG2 KO cell lines using CRISPR-Cas9. Notably, western blot with the UNG antibody showed a complete loss of signal in UNG2 KO cells (Fig. 1H), indicating that the detected band is specific to UNG2. Henceforth, we used this antibody as a UNG2 antibody. As expected, the levels of genomic uracil were higher in UNG2 KO lines than in the parental line (Fig. S1G). Similar to UNG knockdown cells, the UNG2 KO cell lines were more sensitive to ATRi than parental U2OS cells (Fig. 1I, S1H).

In addition to UNG, three other DNA glycosylases, including SMUG1, MBD4, and TDG, have been implicated in the processing of genomic uracil in specific contexts (Visnes et al., 2009). SMUG1 excises 5-hydroxymethyl uracil (5-hmdU) and is a possible backup for UNG for removing genomic uracil (Fugger et al., 2021). MBD4 and TDG are primarily involved in removing the thymine generated by the deamination of 5-methylcytosine, but also remove a variety of other base lesions, including U in G:U pairs (Sjolund et al., 2013). To test whether these glycosylases compensate for the loss of UNG to confer ATRi sensitivity, we co-depleted these enzymes with UNG. Loss of SMUG1 alone slightly increased genomic uracil, but co-depletion of UNG and SMUG1 did not increase genomic uracil more than UNG loss alone (Fig. S2A-B), showing that SMUG1 does not efficiently compensate for UNG loss in suppressing overall genomic uracil levels. Furthermore, knockdown of SMUG1 alone increased ATRi sensitivity, and co-depletion of UNG and SMUG1 increased ATRi sensitivity more than single depletions of UNG and SMUG1 (Fig. S2C), suggesting that the uracil derivatives suppressed by SMUG1, such as 5-hmdU, may also increase ATRi sensitivity. Unlike UNG and SMUG1, neither MBD4 nor TDG affected the overall levels of genomic uracil (Fig. S2D-E, S2G-H). When combined with UNG knockdown, neither MBD4 nor TDG depletion increased genomic uracil levels further (Fig. S2E, S2H). Furthermore, depletion of MBD4 or TDB in UNG knockdown cells did not reduce but increased ATRi sensitivity (Fig. S2F, S2I), suggesting that these glycosylases do not compensate for UNG loss to confer ATRi sensitivity by generating BER intermediates. Finally, we used an *in vitro* biochemical assay to test whether the uracil glycosylase activity in cell extracts was compensated in UNG-depleted cells by other enzymes. Depletion of UNG eliminated the detectable uracil glycosylase activity in cell extracts (Fig. S2J), showing that the loss of UNG activity is not compensated by any other enzymes. Together, these results strongly support the notion that the accumulation of genomic uracil increases ATRi sensitivity in the absence of BER.

### Unprocessed genomic uracil impedes replication forks

To directly test whether genomic uracil affects the progression of replication forks, we used DNA fiber assay to analyze UNG knockdown cells. In cells treated with increasing concentrations of UNG siRNA, UNG was depleted, and replication fork speed was reduced in a manner dependent on siRNA concentrations (Fig. 2A). The slowing of replication forks upon UNG loss was also observed using multiple UNG siRNAs (Fig. 2B, S3A) and UNG2 KO cell lines (Fig. S3B). Thus, UNG promotes replication fork progression in a dose-dependent manner.

**Figure 2.**
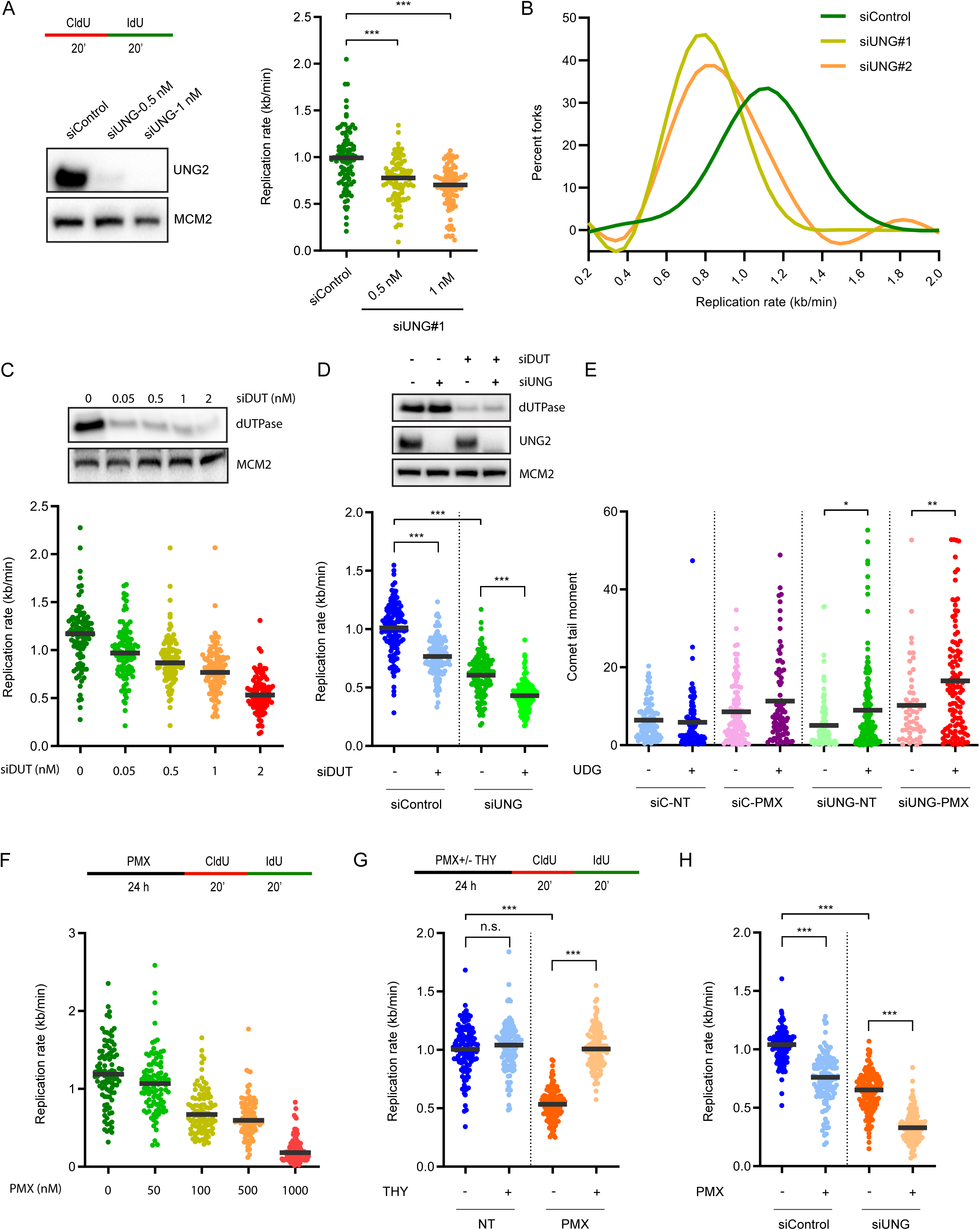
Unprocessed genomic uracil interferes with replication fork progression. (A) U2OS cells were transfected with indicated concentrations of UNG siRNA#1 for 48 h. Levels of UNG2 in whole-cell extracts were analyzed by western blot, and replication rate was analyzed by DNA fiber assay as indicated (n=125 fibers per condition). One of two independent experiments is shown. MCM2 is shown as the loading control in western blot. The asterisk indicates a non-specific band detected by UNG antibody. (B) Graphs show the distribution of replication rate in U2OS cells treated with indicated siRNAs (n=125 fibers per condition). One of two independent experiments is shown. (C) U2OS cells were transfected with indicated concentrations of dUTPase (DUT) siRNA for 48 h. Levels of dUTPase in whole-cell extracts were analyzed by western blot, and replication rate was analyzed by DNA fiber assay as in (A) (n=125 fibers per condition). One of two independent experiments is shown. MCM2 is shown as the loading control in western blot. (D) U2OS cells were transfected with dUTPase (DUT) and UNG#1 siRNAs for 48 h. Levels of dUTPase and UNG2 in whole-cell extracts were analyzed by western blot, and replication rate was analyzed by DNA fiber assay as in (A) (n=125 fibers per condition). One of two independent experiments is shown. MCM2 is shown as the loading control in western blot. (E) U2OS cells transfected with control or UNG#1 siRNA and treated with 200 nM PMX for 24 h were analyzed by U-comet assay. Dot plots represent comet tail moment in dividual cells, and bars display the mean in cell populations (n>50 cells per condition). One of two independent experiments is shown. (F) U2OS cells were treated with indicated concentrations of Pemetrexed (PMX) for 24 h and replication rate was analyzed by DNA fiber assay as indicated (n=125 fibers per condition). One of two independent experiments is shown. (G) U2OS cells were treated with 500 nM PMX and 100 µM Thymidine (THY) for 24 h and replication rate was analyzed by DNA fiber assay as indicated (n=125 fibers per condition). One of two independent experiments is shown. (H) U2OS cells were transfected with control or UNG#1 siRNA for 48 h and treated with 200 nM PMX for 24 h. Replication rate was analyzed by DNA fiber assay as in (F) (n=125 fibers per condition). One of two independent experiments is shown.

Next, we tested whether misincorporation of dUTP to genomic DNA interferes with replication forks. dUTPase (DUT) is an enzyme that hydrolyzes dUTP to dUMP, which in turn serves as a precursor for thymidylate synthesis (Ladner et al., 1996). Depletion of dUTPase is expected to increase the cellular dUTP/dTTP ratio and dUTP misincorporation. Indeed, dUTPase knockdown increased genomic uracil (Fig. S3C). knockdown of dUTPase with increasing concentrations of siRNA reduced fork speed in a dose-dependent manner (Fig. 2C). Importantly, knockdown of dUTPase in cells lacking UNG further reduced fork speed (Fig. 2D), suggesting that dUTP misincorporation induces RS even when genomic uracil is not processed by BER.

To further test whether altered dUTP/dTTP ratio affects fork progression, we sought to inhibit dTTP synthesis. Pemetrexed (PMX) is a folate analogue that inhibits the thymidylate synthase (TYMS), which converts dUTP to dTTP (Shih et al., 1997). Treatment of cells with PMX is expected to reduce dTTP, increase the dUTP/dTTP ratio, and enhance uracil misincorporation. Indeed, as revealed by the U-comet assay, the levels of genomic uracil were increased by PMX treatment (Fig. 2E). Notably, PMX further increased genomic uracil in UNG knockdown cells (Fig. 2E), suggesting that the PMX-induced genomic uracil is removed by BER. In addition, we used UDG to excise uracil from genomic DNA, followed by liquid chromatography-mass spectrometry (LC-MS) to measure uracil levels. LC-MS confirmed that after PMX treatments, UNG2 KO cells contained more genomic uracil than parental U2OS cells (Fig. S3D). Treatment of cells with PMX reduced fork speed in a dose-dependent manner (Fig. 2F). Importantly, the effect of PMX on fork speed was reversed by supplementing cells with a low concentration of thymidine (Fig. 2G), which increases dTTP levels but does not inhibit DNA synthesis by reducing the dCTP pool (Bjursell and Reichard, 1973), confirming that PMX decreases fork speed by reducing dTTP and increasing uracil misincorporation. Treatment of UNG knockdown cells with PMX further reduced fork speed (Fig. 2H), showing that PMX affects fork progression even when genomic uracil is not processed by BER. Together, the experiments using UNG-depleted cells, dUTPase-depleted cells, and PMX-treated cells strongly suggest that unprocessed genomic uracil interferes with replication forks and induces RS.

### Genomic uracil induces PrimPol-generated ssDNA gaps

To understand how genomic uracil impedes replication forks, we asked whether uracil interferes with DNA polymerases in cells. Stalling of DNA polymerases by barriers in DNA typically results in formation of ssDNA ahead of polymerases, allowing the ssDNA-binding protein complex RPA to accumulate adjacent to stalled polymerases (Byun et al., 2005). To test the effects of genomic uracil on DNA polymerases, we performed Proximity Ligase Assay (PLA) with antibodies to the catalytic subunit of DNA polymerase epsilon (POLE1) and the RPA32 subunit of RPA. PLA foci were readily detected in hydroxyurea (HU)-treated cells when both POLE1 and RPA32 antibodies were applied, but not when either antibody was used individually (Fig. S3E), validating this assay for the detection of polymerase stalling. Importantly, PMX treatment significantly increased PLA foci in both control cells and UNG knockdown cells (Fig. 3A, 3B), suggesting that genomic uracil increases POLE stalling independently of UNG-mediated BER.

**Figure 3.**
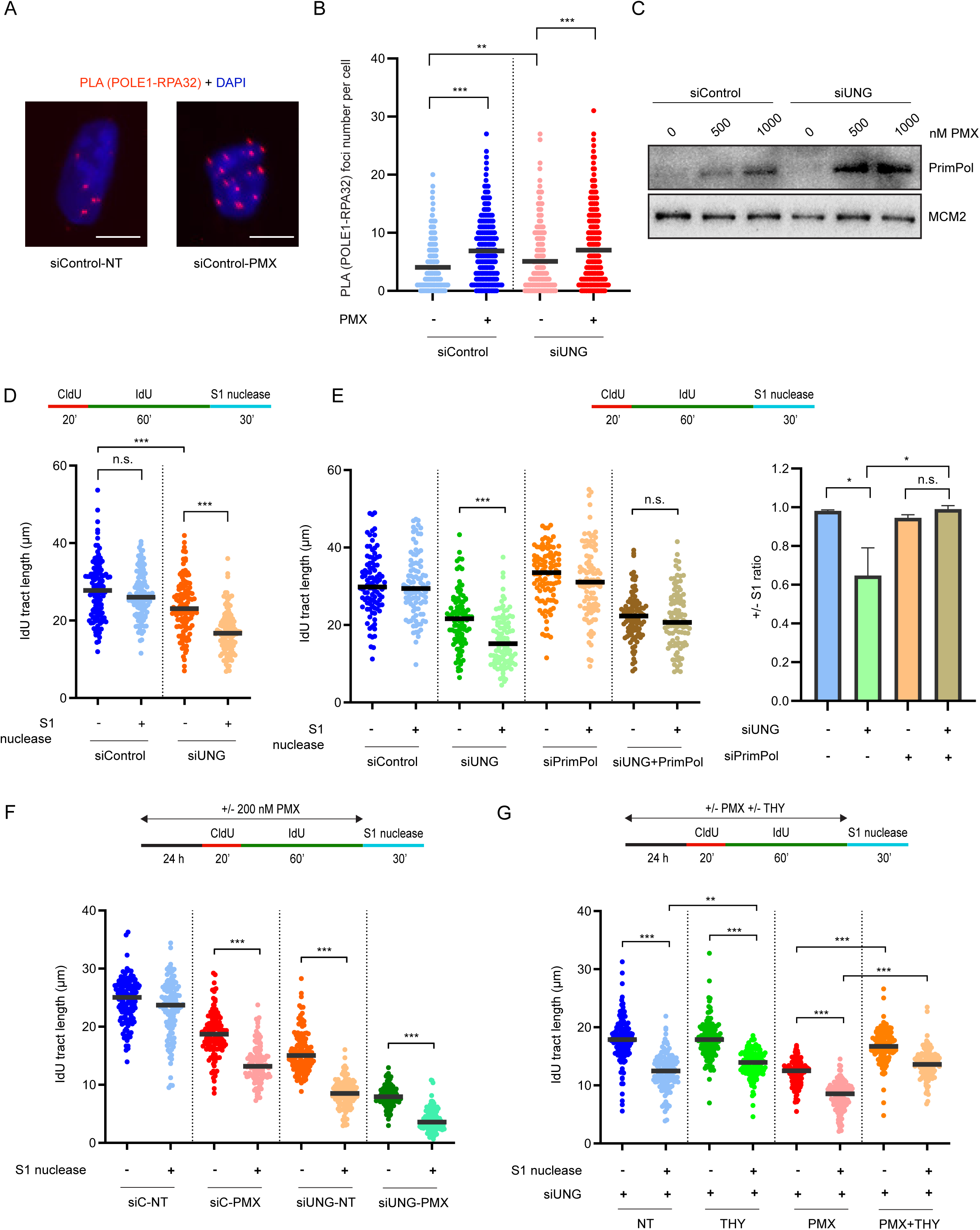
Genomic uracil induces PrimPol-dependent ssDNA gaps in nascent DNA. (A and B) U2OS cells transfected with control or UNG#1 siRNA and treated with 100 nM PMX for 24 h were analyzed by PLA using anti-POLE1 and anti-RPA32 antibodies. Representative images of PLA foci are shown (A) and quantified in individual cells (B) (n>400 cells per condition). One of three independent experiments is shown. Scale bar, 10 µM. (C) U2OS cells were transfected with control or UNG#1 siRNA and treated with indicated concentrations of PMX for 24 h. Chromatin fractions were prepared and blotted for PrimPol. MCM2 is shown as the loading control. (D) U2OS cells were transfected with control or UNG#1 siRNA for 48h, and cells were analyzed by DNA fiber assay as indicated. DNA fibers were treated with S1 nuclease (20 U/mL) for 30 min at 37°C, and the length of IdU replication tracts (n=125 fibers per condition) was measured. One of two independent experiments is shown. (E) U2OS cells were transfected with UNG#1 and PrimPol siRNA for 48 h and analyzed as in (D). The length of IdU replication tracts (n=125 fibers per condition) was measured. One of two independent experiments is shown. Right, ratio of IdU tracts from S1-treated to -untreated fibers. Data are displayed as mean ± s.d. from two independent experiments. (F) U2OS cells transfected with control or UNG#1 siRNA were treated with 200 nM PMX for 24 h and analyzed as in (D). The length of IdU replication tracts (n=125 fibers per condition) was measured. One of two independent experiments is shown. (G) U2OS cells transfected with UNG#1 siRNA were treated with 500 nM PMX and 100 µM thymidine (THY) for 24 h and analyzed as in (D). The length of IdU replication tracts (n=125 fibers per condition) was measured. One of two independent experiments is shown.

DNA polymerase stalling and accumulation of RPA-coated ssDNA at replication forks is known to trigger PrimPol-mediated repriming, which enables resumption of DNA synthesis ahead of stalled polymerases but leaves ssDNA gaps in nascent DNA (Bianchi et al., 2013; Guilliam et al., 2017; Mouron et al., 2013). In cells treated with increasing concentrations of PMX, PrimPol was recruited onto chromatin in a PMX dose-dependent manner (Fig. S3F). Furthermore, PMX further increased the levels of chromatin-bound PrimPol in UNG knockdown cells (Fig. 3C), suggesting that genomic uracil promotes the recruitment of PrimPol to stalled forks independently of UNG-mediated BER.

To directly test whether genomic uracil induces PrimPol-mediated ssDNA gaps, we treated DNA fibers with the S1 nuclease, which specifically cleaves ssDNA (Quinet et al., 2017). If ssDNA gaps are present in nascent DNA, replication tracts should be shortened by S1. Knockdown of UNG significantly increased the shortening of replication tracts by S1 (Fig. 3D, S3G), and similar observations were made in UNG2 KO cell lines (Fig. S3I-J), showing that UNG loss increases ssDNA gaps. Notably, the increase of S1 cleavage of replication tracts in UNG knockdown and UNG2 KO cells was not observed when PrimPol was co-depleted (Fig. 3E, S3H-J), showing that ssDNA gaps were formed in a PrimPol-dependent manner. We noted that loss of PrimPol did not significantly alter fork speed in control, UNG knockdown, and UNG2 KO cells (Fig. 3E, S3J), suggesting that cells use an alternative mechanism to sustain fork progression in the absence of PrimPol.

PMX treatment and dUTPase knockdown also increased the shortening of replication tracts by S1 (Fig. 3F, S3K, S3L), supporting the idea that genomic uracil induces ssDNA gaps. Knockdown of PrimPol prevented the shortening of replication tracts by S1 in PMX-treated cells (Fig. S3K), showing that the ssDNA gaps formed in this context were also PrimPol-dependent. In UNG knockdown cells, replication tracts were further shortened by PMX treatment or dUTPase depletion (Fig. 3F, S3L), reflecting a further increase of genomic uracil. Notably, these short tracts were still efficiently cleaved by S1 (Fig. 3F, S3L), suggesting that ssDNA gaps were formed independently of UNG-mediated BER. Supplementing UNG knockdown cells with thymidine reduced the cleavage of replication tracts by S1 (Fig. 3G, lanes 1-4). In the presence of PMX, thymidine partially rescued fork speed in UNG knockdown cells and also reduced S1 cleavage (Fig. 3G, lanes 5-8). Thus, unprocessed genomic uracil, which is increased by UNG loss and PMX-induced uracil misincorporation, induces PrimPol-generated ssDNA gaps.

### Exogenous dUTP induces replication stress

To test the effects of dUTP misincorporation on replication forks more directly, we sought to directly increase dUTP levels in cells. Because dNTPs do not permeate cell membranes efficiently, we tested whether a recently developed synthetic nucleoside triphosphate transporter (henceforth termed Bio-Tracker, BT) can transport dUTP into cells (Zawada et al., 2018). We first tested BT with Cy3-labeled dUTP and validated the import of Cy3-dUTP into U2OS cells (Fig. 4A). Using DNA fiber assay, we confirmed that BT-imported Cy3-dUTP was incorporated into genomic DNA (Fig. 4A). As expected, our U-comet assay showed a significant increase of genomic uracil in cells treated with BT and Cy3-dUTP (Fig. 4B). These experiments established a strategy to directly increase the dUTP pool in cells and dUTP misincorporation to the genome.

**Figure 4.**
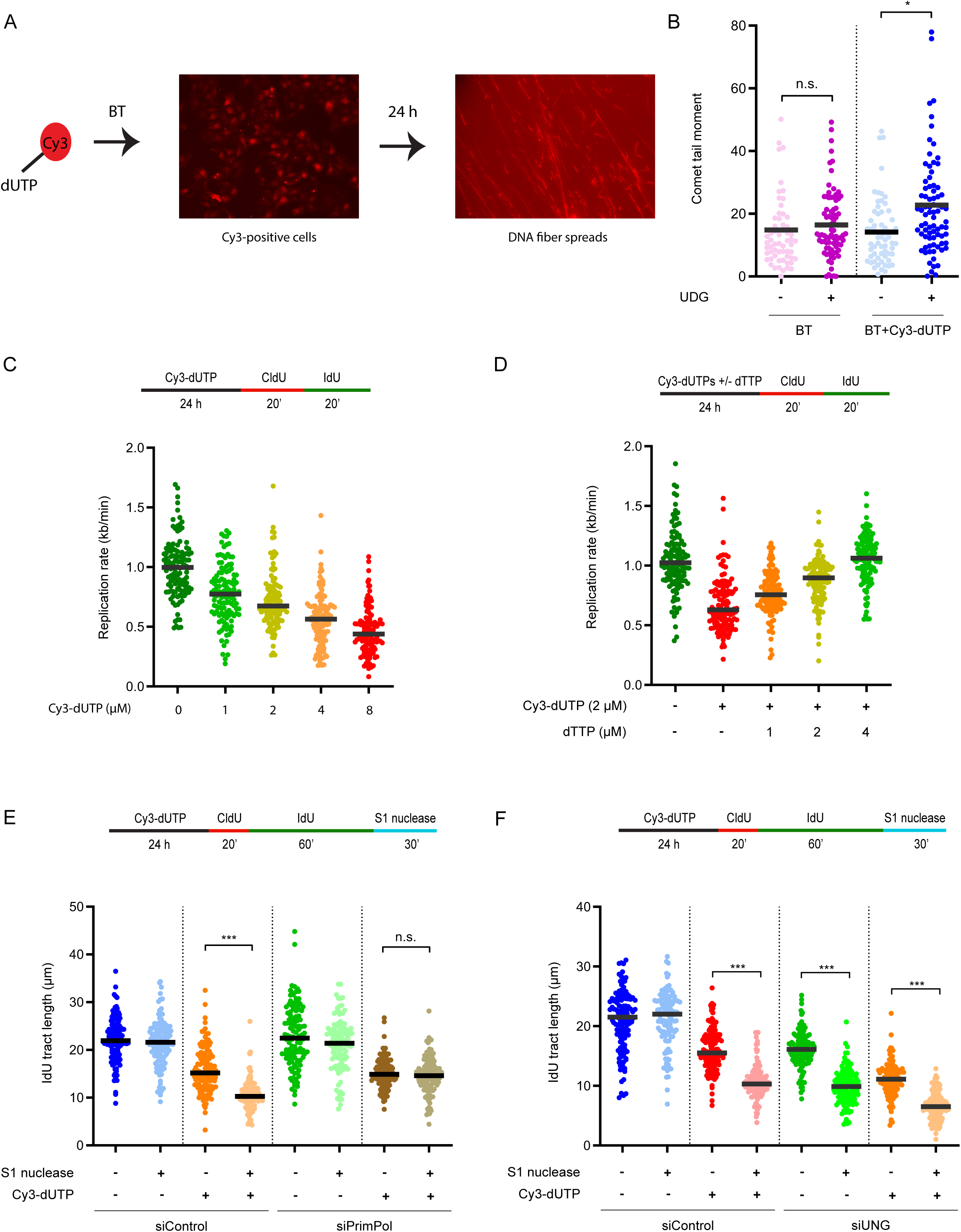
Exogenous dUTP supplementation induces replication stress. (A) A schematic of import of Cy3-dUTP into live cells by Bio-Tracker (BT). U2OS cells were treated with 2 µM BT and 2 µM Cy3-dUTP for 24 h in Leibovitz’s L-15 Medium and analyzed by fluorescence. After 24 h, cells were analyzed by DNA fiber to visualize the incorporation of Cy3-dUTPs to genomic DNA. (B) U2OS cells were treated with 2 µM BT and 2 µM Cy3-dUTP for 24 h and analyzed by U-comet assay. Dot plots represent comet tail moment in individual cells, and bars display the mean in cell populations (n>50 cells per condition). One of two independent experiments is shown. (C) U2OS cells were treated with 2 µM BT and indicated concentrations of Cy3-dUTP for 24 h and replication rate was analyzed by DNA fiber assay as indicated (n=125 fibers per condition). One of two independent experiments is shown. (D) U2OS cells were treated with 2 µM BT, 2 µM Cy3-dUTP and indicated concentrations of dTTP for 24 h, and replication rate was analyzed by DNA fiber assay as indicated (n=125 fibers per condition). One of two independent experiments is shown. (E) U2OS cells transfected with control or PrimPol siRNA were treated with 2 µM BT and 2 µM Cy3-dUTP for 24 h and analyzed by DNA fiber assay as indicated. DNA fibers were treated with S1 nuclease (20 U/mL) for 30 min at 37°C, and the length of IdU replication tracts (n=125 fibers per condition) was measured. One of two independent experiments is shown. (F) U2OS cells were transfected with control or UNG#1 siRNA, treated with 2 µM BT and 2 µM Cy3-dUTP for 24 h, and analyzed as in (E).

Next, we sought to characterize the cellular response to genomic uracil using this system. Import of Cy3-dUTP reduced fork speed in a dose-dependent manner (Fig. 4C). Importantly, when increasing concentrations of dTTP were co-imported with Cy3-dUTP, the Cy3-dUTP-induced fork slowing was alleviated by dTTP in a dose-dependent manner (Fig. 4D). Similar observations were made using unlabeled dUTP and dTTP (Fig. S4A), ruling out unexpected effects of Cy3. Furthermore, dTTP also suppressed the increase of genomic uracil induced by Cy3-dUTP in a concentration-dependent manner (Fig. S4B). These results strongly suggest that the dUTP/dTTP ratio is a key determinant of genomic uracil levels.

Using DNA fiber assay and the S1 nuclease, we confirmed that both Cy3-dUTP and unlabeled dUTP, but not dTTP, induced ssDNA gaps in nascent DNA (Fig. 4E, S4C). Furthermore, the induction of ssDNA gaps by Cy3-dUTP was dependent on PrimPol (Fig. 4E, S4D). Notably, even in UNG knockdown cells, Cy3-dUTP further shortened replication tracts, and the short tracts were still sensitive to S1 cleavage (Fig. 4F), suggesting that unprocessed genomic uracil impedes replication forks and induces ssDNA gaps. At a low concentration that did not change the ATRi sensitivity of cells treated with control siRNA, Cy3-dUTP increased the ATRi sensitivity of UNG knockdown cells (Fig. S4E), supporting the notion that dUTP misincorporation increases unprocessed genomic uracil and associated RS.

### ATR is required for the repair of uracil-induced ssDNA gaps

Next, we investigated how genomic uracil enhances ATRi sensitivity. PrimPol-generated ssDNA gaps, if not repaired properly, can persist into the next cell cycle, collide with replication forks, and give rise to DSBs (Simoneau et al., 2021; Taglialatela et al., 2021; Tirman et al., 2021). To test whether ATR is involved in the repair of uracil-induced ssDNA gaps, we pulse-labeled nascent DNA in UNG knockdown cells and then let cells progress through the cell cycle in the presence or absence of ATRi (Fig. 5A). If ssDNA gaps are formed during DNA labeling and subsequently repaired, ssDNA gaps should be gradually removed from labeled DNA over time. Using the S1 nuclease, we confirmed the increase of ssDNA gaps in UNG knockdown cells right after nascent DNA labeling (Fig. 5A, lanes 1-4). In the absence of ATRi, ssDNA gaps gradually disappeared from labeled DNA after 8 and 24 h (Fig. 5A, lanes 5-6), showing repair of the gaps. In the presence of ATRi, however, the removal of ssDNA gaps was significantly compromised (Fig. 5A, lanes 7-8). Thus, ATR is required for the efficient repair of uracil-induced ssDNA gaps.

**Figure 5.**
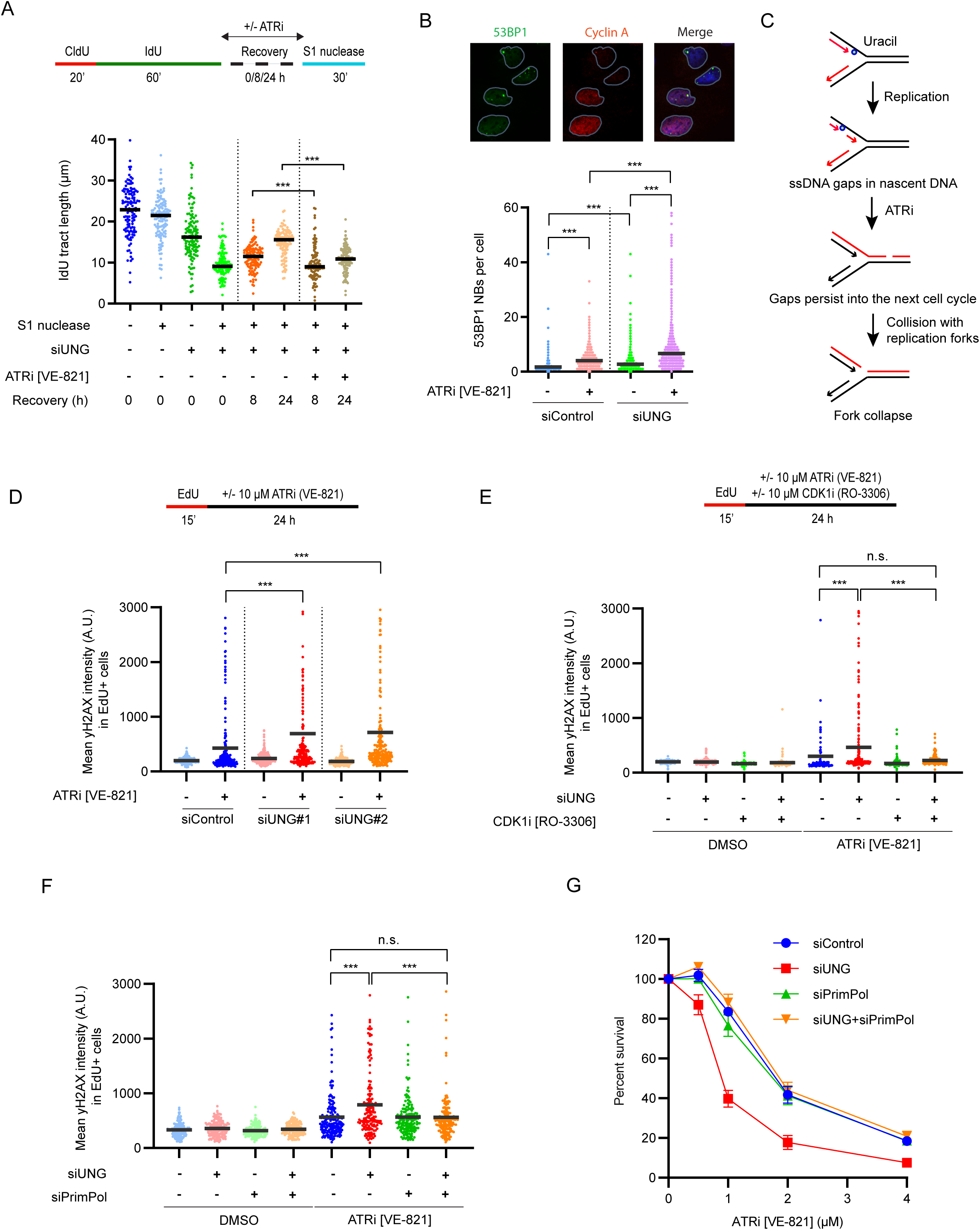
ATRi prevents repair of uracil-induced ssDNA gaps and increases DNA damage. (A) S1 nuclease assay to determine the kinetics of ssDNA gap repair. U2OS cells were transfected with control or UNG#1 siRNA, sequentially labeled with CldU and IdU and cultured in the presence of absence of ATRi for 0, 8 or 24 h, as indicated. At each time point, cells were analyzed by DNA fiber assay. DNA fibers were treated with S1 nuclease (20 U/mL) for 30 min at 37°C, and the length of IdU replication tracts (n=125 fibers per condition) was measured. One of two independent experiments is shown. (B) U2OS cells transfected with control or UNG#1 siRNA and treated with 2 µM ATRi (VE-821) for 48 h. Cyclin A-negative G1 cells that contained 53BP1 nuclear bodies (NBs) were quantified (n>1200 cells per condition). One of two independent experiments is shown. Representative images are shown. (C) Model for the generation of DNA damage by ATRi in cells lacking UNG2. In the absence of UNG2, cells accumulate high levels of uracil in the genome, resulting in PrimPol-mediated ssDNA gaps in the newly replicated DNA (shown in red). Upon ATR inhibition, these gaps remain unrepaired and persist into the next cell cycle. In the next S phase, encounter of ongoing replication forks with these gaps would cause fork collapse and DSB generation. (D) U2OS cells were transfected with indicated siRNAs and pulse labeled with 5 µM EdU for 15 min, followed by treatment with 10 µM ATRi (VE-821) for 24 h. gH2AX intensity in EdU-positive cells was quantified (n>250 cells per condition). One of two independent experiments is shown. (E) U2OS cells were transfected with UNG#1 and pulse labeled with 5 µM EdU for 15 min, followed by treatment with 10 µM ATRi (VE-821) +/-10 µM CDK1i (RO-3306) for 24 h. gH2AX intensity in EdU-positive cells was quantified (n>250 cells per condition). One of three independent experiments is shown. (F) U2OS cells were transfected with UNG#1 and PrimPol siRNA and pulse labeled with 5 µM EdU for 15 min, followed by treatment with 10 µM ATRi (VE-821) for 24 h. gH2AX intensity in EdU-positive cells was quantified (n>250 cells per condition). One of three independent experiments is shown. (G) U2OS cells were transfected with UNG#1 and PrimPol siRNA and treated with indicated concentrations of ATRi (VE-821) for 5-7 days. Cell viability was determined using CellTiter-Glo and normalized to the cells untreated with ATRi under each condition. Data are shown as mean ± s.d. (n=2 independent experiments).

If ATRi prevents the repair of uracil-induced ssDNA gaps, these gaps may persist into the next cell cycle. To test this possibility, we analyzed whether the 53BP1 nuclear bodies (NBs) in G1 phase are affected by ATRi in UNG knockdown cells. G1 53BP1 NBs are induced by RS and associated with under-replicated DNA (Harrigan et al., 2011; Lukas et al., 2011). In the absence of ATRi, 53BP1 NB levels in Cyclin A-negative G1 cells were higher in the UNG knockdown cell population than in the control cell population (Fig. 5B), consistent with the induction of RS by UNG loss. In the presence of ATRi, 53BP1 NBs were further increased in UNG knockdown cells (Fig. 5B), suggesting that more under-replicated DNA persisted into the next G1. These results raised the possibility that uracil-induced ssDNA gaps can persist into the next S phase in the presence of ATRi, leading to replication fork collapse (Simoneau et al., 2021) (Fig. 5C). To test this idea, we pulse labelled S-phase cells with EdU and then treated them with ATRi for 24 h to analyze the effects of ATR in these cells in the second cell cycle. ATRi-induced γH2AX was significantly higher in EdU^+^ UNG knockdown cells than in EdU^+^ control cells (Fig. 5D), suggesting that genomic uracil promotes ATRi-induced DSB formation. Furthermore, blocking the cell cycle at the G2/M transition with the CDK1 inhibitor R0-3306 drastically reduced ATRi-induced γH2AX (Fig. 5E), showing that the entry to the next cell cycle is required for ATRi-induced DSB formation. Importantly, knockdown of PrimPol reversed the induction of γH2AX in the UNG-depleted cells (Fig. 5F), confirming that the DSBs arose from PrimPol-generated ssDNA gaps. Remarkably, co-depletion of PrimPol in UNG knockdown cells completely reversed the increase of ATRi sensitivity (Fig. 5G, S5A), lending further support to the notion that the ATRi sensitivity of UNG knockdown cells is attributed to PrimPol-generated ssDNA gaps. Together, these results suggest that ATRi induces gap-derived DSBs in UNG-depleted cells in a trans-cell-cycle manner.

To understand how replication forks collapse in ATRi-treated UNG knockdown cells, we asked whether nucleases are involved. The DNA structure-specific nucleases MUS81 and MRE11 have been implicated in the collapse of replication forks (Mann et al., 2022; Regairaz et al., 2011). Knockdown of MUS81 and inhibition of MRE11 with Mirin both significantly suppressed ATRi-induced γH2AX in UNG knockdown cells in the second S phase (Fig. S5B-C), suggesting that both of these nucleases are involved in the accumulation of ATRi-induced DSBs in UNG-depleted cells. It is possible that when the repair of uracil-induced gaps is compromised by ATRi, these gaps or the forks approaching them need to be processed by MRE11 and MUS81 to form DSBs efficiently.

### High UNG expression in cancer cells is associated with uracil-induced replication stress

The role of UNG in limiting genomic uracil and RS raises the possibility that UNG may be upregulated in a subset of cancers to allow cancer cells to cope with uracil-induced RS. Notably, *UNG2* is commonly expressed at higher levels in lung cancer cell lines compared to non-malignant lung epithelial cells (Weeks et al., 2013). To test whether cancer cells expressing high levels of UNG2 depend on it to suppress genomic uracil and RS, we analyzed UNG2 protein levels in a panel of lung cancer cell lines (Fig. 6A). Among the cell lines, only H1299 expresses UNG2 at high levels. No correlation between UNG2 protein levels and ATRi sensitivity was observed in these cell lines (Fig. S6A). However, when UNG was knocked down in these cell lines, H1299, but not the other cell lines, became significantly more sensitive to ATRi (Fig. 6B, S6B). These results suggest that H1299, which expresses UNG2 at the highest level in this cell line panel, is the most dependent on UNG2 to suppress RS. Similar to the cancer cell lines expressing low levels of UNG2, RPE-1, a non-transformed cell line, expressed UNG2 at a low level and exhibited only a modest increase in ATRi sensitivity upon UNG knockdown (Fig. S6C).

**Figure 6.**
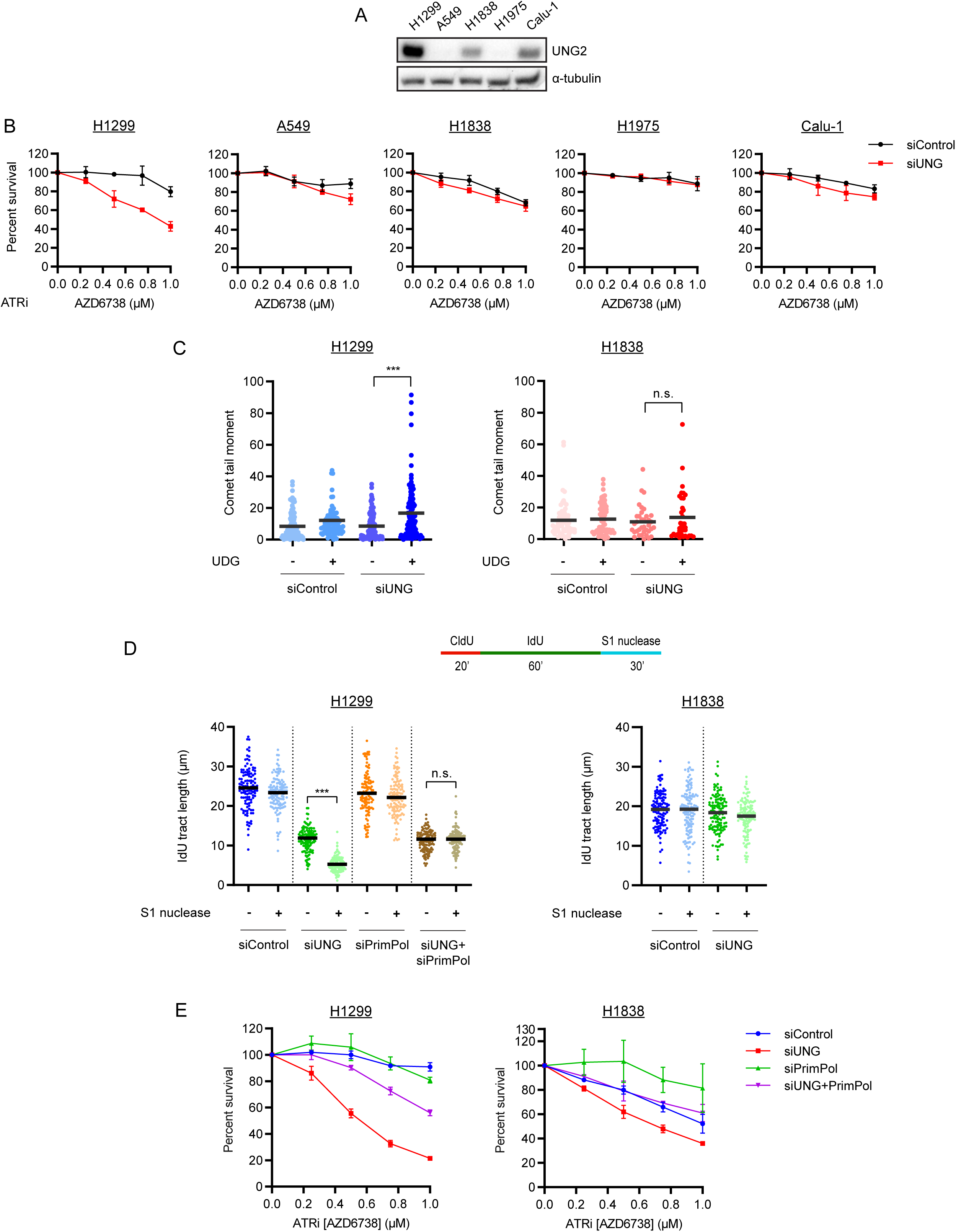
A subset of cancer cells rely on UNG2 to suppress replication stress. (A) Levels of UNG2 in whole-cell extracts of indicated cells were analyzed by western blot. α-tubulin is shown as the loading control. (B) Indicated cell lines were transfected with control or UNG#1 siRNA and treated with indicated concentrations of ATRi (AZD6738) for 5-7 days. Cell viability was determined using CellTiter-Glo and normalized to the cells untreated with ATRi under each condition. Data are shown as mean ± s.d. (n=4 independent experiments). (C) H1299 and H1838 cells transfected with control or UNG#1 siRNA were analyzed by U-comet assay. Dot plots represent comet tail moment per cell and bars display the mean (n>50 cells per condition). One of two independent experiments is shown. (D) H1299 and H1838 cells were transfected with control or UNG#1 and PrimPol siRNAs for 48 h, and cells were analyzed by DNA fiber assay as indicated. DNA fibers were treated with S1 nuclease (20 U/mL) for 30 min at 37°C, and the length of IdU replication tracts (n=125 fibers per condition) was measured. One of two independent experiments is shown. (E) H1299 and H1838 cells were transfected with UNG#1 and PrimPol siRNA and treated with indicated concentrations of ATRi (AZD6738) for 5-7 days. Cell viability was determined using CellTiter-Glo and normalized to the cells untreated with ATRi under each condition. Data are shown as mean ± s.d. (n=2 independent experiments).

To directly compare the levels of genomic uracil in the lung cancer cell lines, we analyzed H1299 and H1838, which expresses low levels of UNG2 (Fig. 6A), with U-comet assay. Upon UNG knockdown, a drastic increase of genomic uracil was detected in H1299 but not H1838 (Fig. 6C). These results demonstrate that H1299 is more dependent on UNG2 than H1838 to suppress genomic uracil, providing an example of the UNG2 dependency of cancer cells.

To characterize the RS in cancer cells expressing high or low levels of UNG2, we analyzed H1299 and H1838 with DNA fiber assay. Knockdown of UNG drastically reduced replication fork speed in H1299 cells and increased ssDNA gaps in a PrimPol-dependent manner (Fig. 6D, S6D-E). In contrast, UNG loss did not affect fork speed and only slightly increased ssDNA gaps in H1838 cells (Fig. 6D, S6D). Furthermore, in H1299, PrimPol knockdown significantly reversed the increase of ATRi sensitivity induced by UNG loss (Fig. 6E). In H1838, PrimPol knockdown modestly increased ATRi resistance and also completely reversed the modest increase of ATRi sensitivity induced by UNG depletion (Fig. 6E). Thus, in both H1299 and H1838, uracil-induced RS and ATRi sensitivity are largely attributed to PrimPol-generated ssDNA gaps. H1299, which is more dependent on UNG2 to suppress genomic uracil, is also more sensitive to ATRi upon UNG loss.

### Induction of genomic uracil sensitizes UNG-dependent cancer cells to ATRi

Given our finding that some cancer cells are dependent on UNG2 to suppress genomic uracil and RS, we asked whether we could sensitize these cancer cells to ATRi by increasing genomic uracil without depleting UNG2. First, we knocked down dUTPase in H1299 and H1838 and tested the effects on ATRi sensitivity (Fig. S7A). dUTPase depletion increased ATRi sensitivity drastically in H1299 but only modestly in H1838. Next, we used PMX to increase genomic uracil in H1299 and H1838 cells. Remarkably, PMX significantly increased the ATRi sensitivity of H1299 cells but not H1838 cells (Fig. 7A, S7B). Furthermore, PMX also drastically increased the ATRi sensitivity of HL-60, an AML cell line expressing UNG at high levels (Fig. S7B) (Human Protein Atlas proteinatlas.org). Importantly, the PMX-induced ATRi hypersensitivity of H1299 cells was completely reversed by thymidine (Fig. 7B), confirming that PMX enhances ATRi sensitivity by reducing dTTP and increasing the dUTP/dTTP ratio. Together, these results suggest that PMX can be used to sensitize UNG-dependent cancer cells to ATRi.

**Figure 7.**
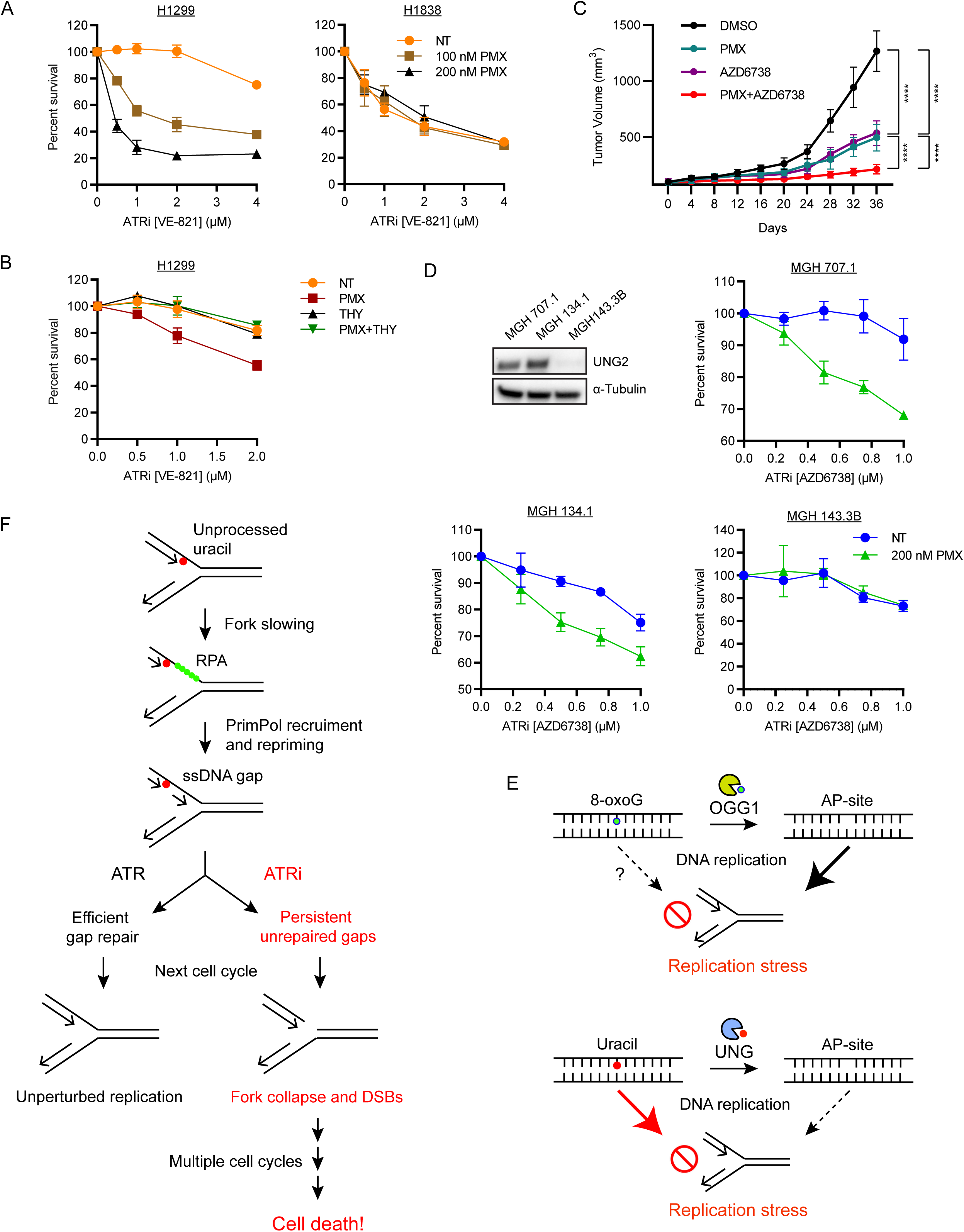
Induction of genomic uracil sensitizes UNG-dependent cancer cells to ATRi. (A) H1299 and H1838 cells were treated with indicated concentrations of PMX and ATRi (VE-821) for 5-7 days. Cell viability was determined using CellTiter-Glo and normalized to the cells untreated with ATRi under each condition. Data are shown as mean ± s.d. (n=2 independent experiments). (B) H1299 cells were treated with 100 nM PMX, 50 µM THY and indicated concentrations of ATRi (VE-821) for 5-7 days. Cell viability was determined using CellTiter-Glo and normalized to the cells untreated with ATRi under each condition. Data are shown as mean ± s.d. (n=2 independent experiments). (C) H1299 xenograft model. Growth curves of H1299 tumors in control mice and mice treated with 50 mg/kg ATRi (AZD6738) and/or 75 mg/kg PMX. n=4 mice in each group. (D) MGH 707.1, MGH 134.1 and MGH 143-3B cells were treated with 200 nM PMX and indicated concentrations of ATRi (AZD6738) for 5-7 days. Cell viability was determined using CellTiter-Glo and normalized to the cells untreated with ATRi under each condition. Data are shown as mean ± s.d. (n=2 independent experiments). Levels of UNG2 in whole-cell extracts of indicated cells were analyzed by western blot. α-tubulin is shown as the loading control. (E) Model for the induction of replication stress by different types of base lesions. (Top) Some base alterations (e.g., 8-oxoG) induce replication stress by generating BER intermediates. (Bottom) Genomic uracil can induce replication stress through BER-independent and -dependent mechanisms. The relative contributions of these two mechanisms may be determined by the levels of genomic uracil and the efficiency and kinetics of BER. (F) Model for the induction of replication stress by unprocessed uracil. Encounter of replication forks with uracil in the template DNA causes polymerase slowdown, resulting in the accumulation of RPA-coated ssDNA at the fork. The accumulation of RPA-ssDNA leads to the recruitment of PrimPol and repriming ahead of the uracil, generating ssDNA gaps in nascent DNA. Upon ATR inhibition, PrimPol-generated gaps remain unrepaired and persist into the next cell cycle, where they collide with replication forks and generate DSBs. Over multiple cell cycles, ATRi allows genomic uracil to drive accumulation of DNA damage and ultimately cell death.

Next, we asked whether the combination of ATRi and PMX is effective on UNG2-dependent tumors *in vivo*. We used H1299 cells to generate xenograft tumors in female NOD SCID gamma (NSG) mice. Treatment with the ATRi AZD6738 or PMX alone delayed tumor growth as compared with DMSO controls (Fig. 7C). Notably, a significantly higher anti-tumor effect was observed when ATRi and PMX were combined (Fig. 7C and S7C). Neither ATRi nor PMX significantly reduced animal weights at the doses used (Fig. S7D). Thus, the combination of ATRi and PMX is more effective than either drug alone in inhibiting the growth of tumors under uracil-induced RS *in vivo*.

Finally, we asked whether the findings from established lung cancer cell lines can be applied to patient-derived materials. We tested three cell lines derived from lung cancer patients at Massachusetts General Hospital (MGH). Among these lines, MGH707.1 and MGH134.1 expressed high levels of UNG2 protein, whereas UNG2 was undetected in MGH143.3B (Fig. 7D). Consistent with the results from H1299 and H1838, knockdown of UNG significantly increased genomic uracil in MGH707.1 and MGH134.1, but not in MGH143.3B (Fig. S7E-F). Furthermore, UNG knockdown significantly reduced fork speed and increased ssDNA gaps in MGH707.1 and MGH134.1 (Fig. S7G). In contrast, in MGH143.3B, UNG depletion only slightly reduced fork speed and did not significantly increase ssDNA gaps (Fig. S7G). Importantly, PMX significantly sensitized MGH707.1 and MGH134.1, but not MGH143.3B, to ATRi (Fig. 7D), showing that PMX effectively sensitizes patient-derived, UNG2-dependent tumor cells to ATRi.

## Discussion

Base alterations in the genome have been long appreciated to be a source of genomic instability. The increase of oxidative stress in cancer cells contributes to tumorigenesis in a variety of oncogenic contexts (Hayes et al., 2020). The induction of BER intermediates by oxidized bases (e.g., 8-oxoG) is believed to increase RS and ultimately give rise to DSBs (Mohni et al., 2019). Indeed, our experiments using OGG1 inhibitor support this model (Fig. 7E). Surprisingly, however, in contrast to this prevailing model, we also found that genomic uracil, a distinct type of base alteration, induces RS independently of UNG-mediated BER. In addition to UNG, we eliminated all known uracil glycosylases as backups for UNG to generate RS through BER intermediates, strengthening the notion that genomic uracil can induce RS independently of BER. In the absence of UNG2, the primary glycosylase that recognizes genomic uracil and initiates BER, replication forks slow down, and cells become increasingly sensitive to ATRi. These results suggest that unprocessed uracil in the genome impedes replication forks even more than the BER intermediates that it generates (Fig. 7E). While genomic uracil interferes with replication forks, our results do not exclude the possibility that uracil-induced BER intermediates also induce RS. The overall impact of genomic uracil on the genome is likely determined by the levels of uracil in DNA and the efficiency and kinetics of BER. If genomic uracil is abundant but does not trigger BER in a timely manner, unprocessed uracil is probably the predominant source of RS. In contrast, if BER is initiated but not completed properly, genomic uracil may induce RS through BER intermediates. It is conceivable that genomic uracil can induce RS in both BER-independent and -dependent manners, and the contributions of these mechanisms may vary in cancer cells depending on the levels of genomic uracil and status of the BER pathway. Notably, recent studies showed that loss of SMUG1 reduced the sensitivity of BRCA1-deficient cells to PARPi (Fugger et al., 2021; Taglialatela et al., 2021), suggesting that AP sites are more toxic than unprocessed hmdU in the context of BRCA1 deficiency and PARP trapping. This finding provides an example of how the genetic background of cancer cells and the action of cancer drugs may influence the effects of uracil and its derivatives in the genome.

In this study, we unexpectedly found that genomic uracil slows down replication forks. This slowdown of replication forks is independent of UNG, suggesting that unprocessed genomic uracil is responsible. Importantly, our PLA analysis using POLE1 and RPA32 antibodies suggests that POLE is uncoupled from the unwinding helicase by genomic uracil in cells, which is a predicted effect of polymerase slowdown. Thus, although previous *in vitro* biochemical studies did not detect a slowdown of DNA polymerases on dU-containing DNA templates (Wardle et al., 2008), genomic uracil impedes DNA polymerases in cells when replication proteins and dNTPs are present in more physiological relevant concentrations. Our experiments also reveal that genomic uracil increases PrimPol-mediated repriming at replication forks, an event triggered by the accumulation of ssDNA and RPA ahead of DNA polymerases. While PrimPol-mediated repriming allows replication forks to progress through the uracil in template, it generates ssDNA gaps that can subsequently give rise to DSBs. Notably, ATRi prevents the efficient repair of uracil-induced gaps. The inhibition of gap repair by ATRi makes these gaps more persistent, leading to fork collapse in the next S phase (Fig. 7F). ATRi also inhibits the repair of collapsed forks, making cancer cells under uracil stress even more dependent on ATR for survival. The trapping of PARP1/2 by PARPi also prevents efficient gap repair (Cong et al., 2021; Simoneau et al., 2021; Vaitsiankova et al., 2022). Furthermore, recent studies implicated both trans-lesion synthesis (TLS) and POLQ in the repair of ssDNA gaps (Belan et al., 2022; Mann et al., 2022; Schrempf et al., 2022; Tirman et al., 2021). It is possible that inhibitors of PARP1/2, the TLS factor REV1, and POLQ can preferentially kill cancer cells under high levels of uracil-associated RS.

What types of cancer cells harbor high levels of genomic uracil? The balance of dUTP and dTTP in cancer cells is likely a key determinant of the frequency of uracil misincorporation and genomic uracil levels (Berger et al., 2008). Indeed, PMX increases the dUTP/dTTP ratio and elevates genomic uracil levels in cancer cells (Bulgar et al., 2012). It is possible that alternations of a number of metabolic pathways in cancer cells increase the dUTP/dTTP ratio and promote the accumulation of uracil in the genome. Furthermore, the cytidine deamination by AID or APOBEC is likely another source of genomic uracil in cancer cells (Petljak and Maciejowski, 2020; Pettersen et al., 2015). Of note, our unpublished data show that expression of APOBEC3A also induces PrimPol-generated ssDNA gaps, which confers ATRi sensitivity. When cancer cells harbor high levels of genomic uracil, they have to cope with uracil-induced RS by limiting genomic uracil and activating the RS response. Both upregulation of BER and metabolic changes reducing the dUTP/dTTP ratio could be used by cancer cells to limit genomic uracil. For example, human lung cancer cells upregulate *UNG* in response to PMX exposure (Weeks et al., 2013). Cancer cells also use TMPK to prevent dUTP incorporation during DNA repair (Hu et al., 2012). We found that cancer cells expressing high levels of UNG2 are more dependent on UNG to suppress genomic uracil. This addiction of cancer cells to suppressors of genomic uracil creates an opportunity to exploit uracil-induced RS. The leukemia cell line HL-60, which expresses high levels of *UNG*, is sensitive to MTHFD2 inhibitors, which reduce thymidine synthesis and increase genomic uracil (Bonagas et al., 2022). Drugs such as PMX or dUTPase inhibitors can also be used to exacerbate the uracil-induced RS in cancer cells. Furthermore, when uracil-induced RS is exacerbated in cancer cells, they become even more dependent on ATR-mediated gap repair and RS response for survival. This model provides a rationale to target cancer cells under high uracil stress using combinations of ATRi with PMX, dUTPase inhibitors, or MTHFD2 inhibitors. In future studies, it is important to identify biomarkers for tumors harboring high uracil stress and develop therapeutics to effectively exacerbate and exploit the uracil-induced RS.

## Acknowledgements

We thank members of Zou, Dyson, Lan, Mostoslavsky, Elia, and Motamedi labs for helpful discussions, Dr. M. Lawrence of bioinformatic analysis, and Dr. C. Ott and the MGH Center for Molecular Therapeutics for cell lines. We thank Meenakshi Basu of the Oh lab for help with RT-qPCR. S.S. was supported by the James A. Harting Scientific Scholar Award from the Rivkin Center. L.Z. is the James and Patricia Poitras Endowed Chair in Cancer Research. This work is supported by the NIH grants CA248526 and CA263934 to L.Z. and funding from the Lungstrong Foundation to A.N.H.

## Author contributions

S.S. and L.Z. designed the study with inputs from other authors. S.S. performed the majority of the experiments and data analysis. P.P. performed the mouse experiment. A.S.K. performed the biochemical assay on uracil glycosylase activity. L.Z. supervised the study. C.S.N., C.R.C. and M.G.V. contributed to the development of a key assay. A.N.H. provided critical reagents. S.S and L.Z. wrote the manuscript with inputs from all authors.

## STAR METHODS

Detailed methods are provided in the online version of this paper and include the following:

- **KEY RESOURCES TABLE**
- **CONTACT FOR REAGENT AND RESOURCE SHARING**
- **EXPERIMENTAL MODEL AND SUBJECT DETAILS**
  - Cell culture
- **METHOD DETAILS**
  - Cloning
  - CRISPR-Cas9 KO cell lines
  - Plasmid transfection
  - RNA interference
  - Quantitative real-time PCR
  - Measurement of AP-sites
  - U-comet assay
  - DNA fiber analysis
  - Uracil excision activity assay
  - Import of dNTPs into live cells
  - Immunofluorescence
  - Proximity ligation assay (PLA)
  - Immunoblots
  - Chromatin fractionation
  - Cell viability assay
  - *In vivo* drug response
- **QUANTIFICATION AND STATISTICAL ANALYSIS**

## KEY RESOURCES TABLE

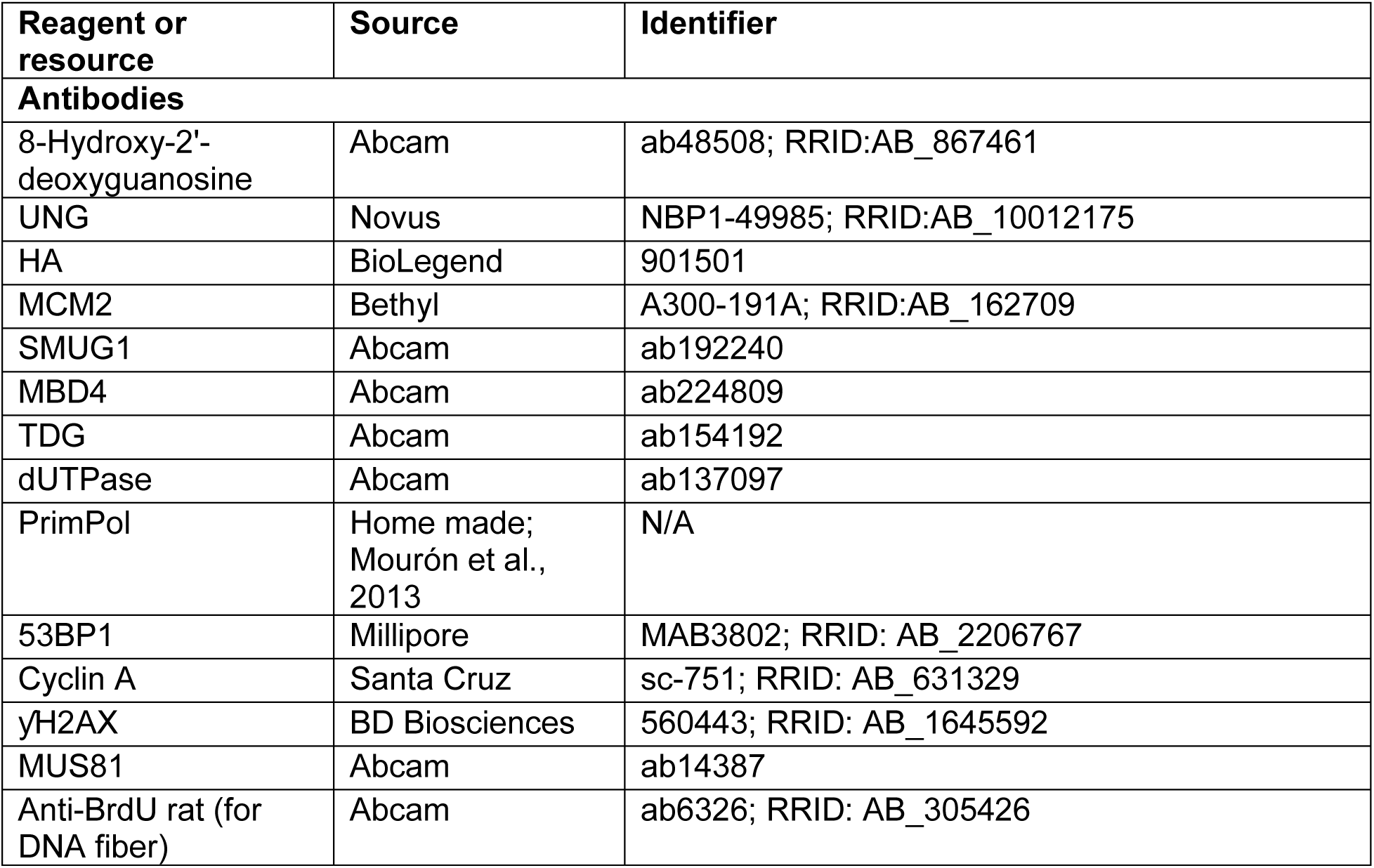
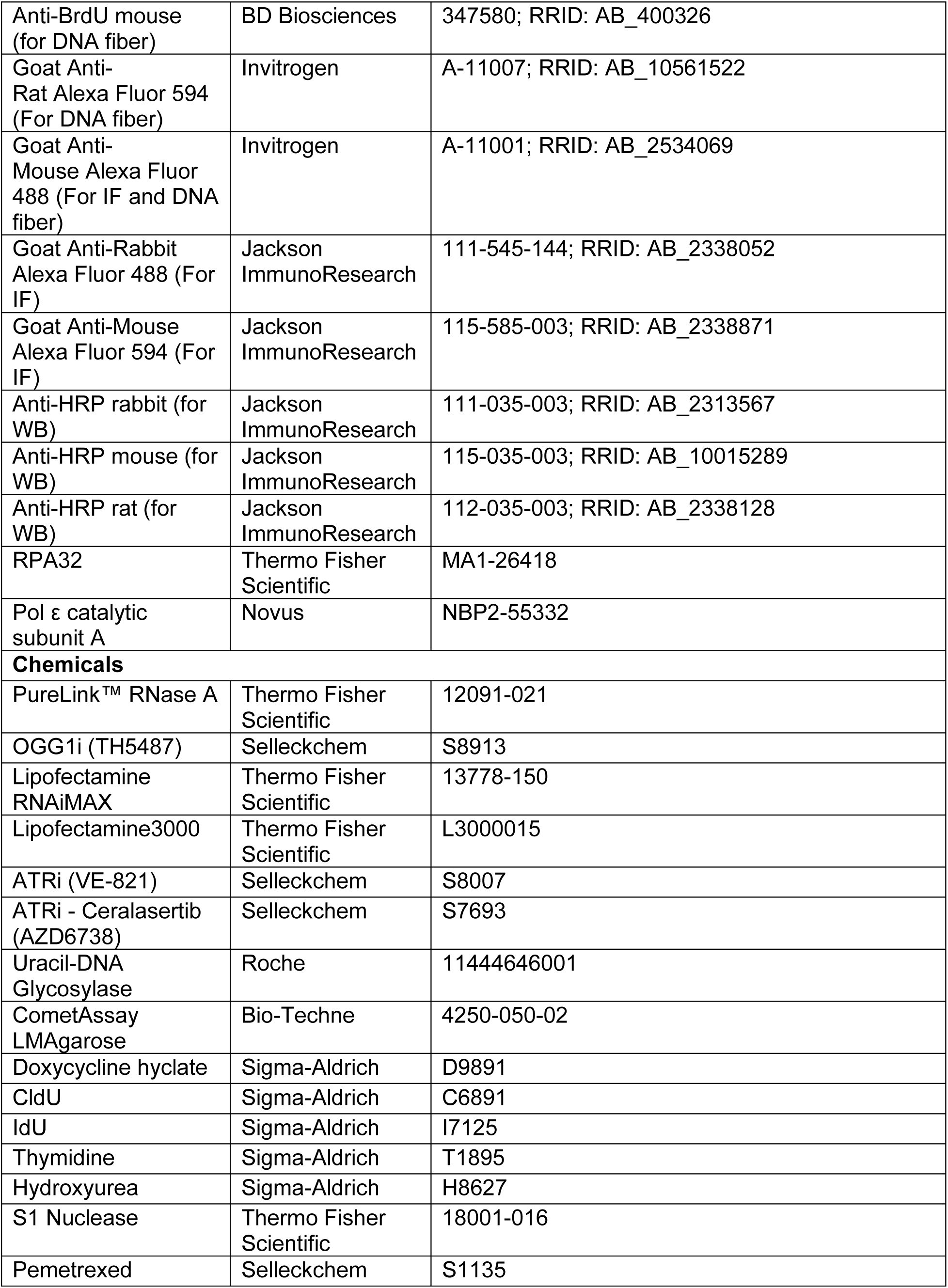
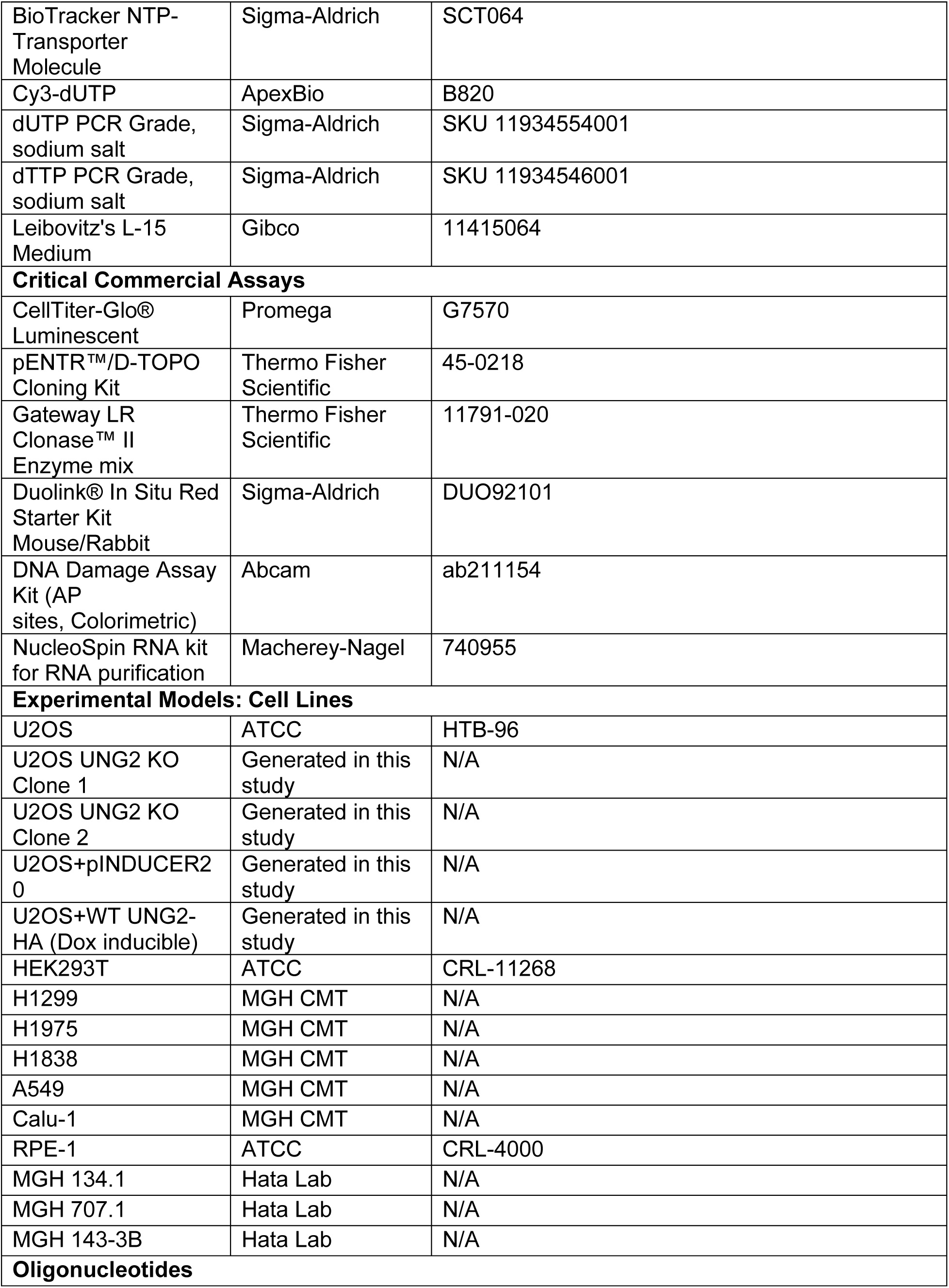
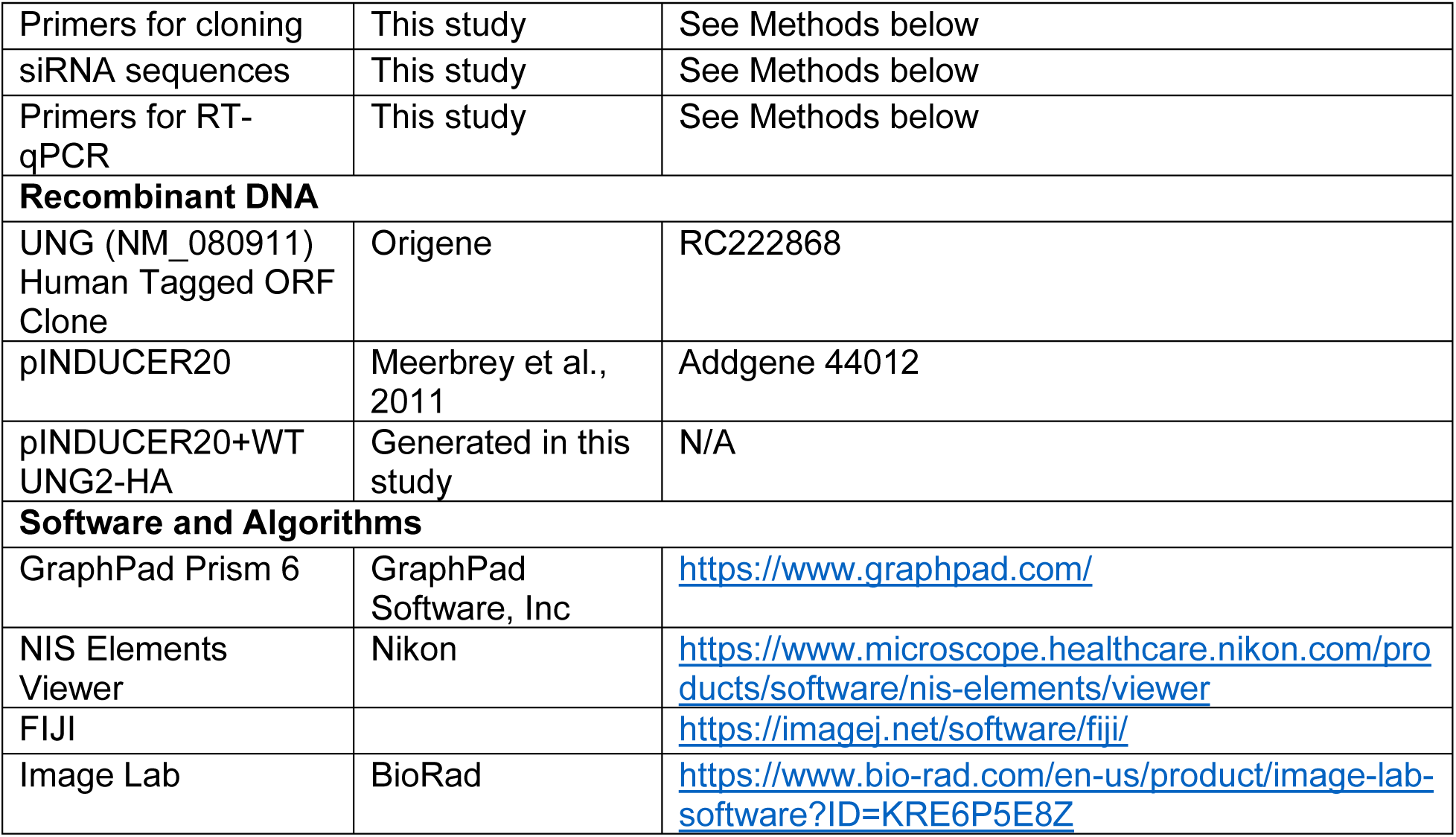

## CONTACT FOR REAGENT AND RESOURCE SHARING

Please direct any requests for further information or reagents to the lead contact, Professor Lee Zou, Department of Pharmacology and Cancer Biology, Duke University School of Medicine, Durham, NC 27710.

## EXPERIMENTAL MODEL AND SUBJECT DETAILS

### Cell culture

The human embryonic kidney cell line HEK293T, the hTERT-immortalized human retinal pigment epithelial cell line RPE-1, the human osteosarcoma cell line U2OS and Its derivative UNG2 KO cell lines were maintained in Dulbecco’s modified Eagle’s medium (DMEM) supplemented with L-glutamine, 10% Fetal Bovine Serum (FBS), and 1% penicillin/streptomycin (PS). Human lung cancer cell lines H1299, H1838, and H1975 were maintained in RPMI 1640 medium supplemented with 10% FBS and 1% PS. The human lung cancer cell line A549 was maintained in F-12K medium supplemented with 10% FBS and 1% PS. The human lung cancer cell line Calu-1 was maintained in McCoy’s 5A medium supplemented with 10% FBS and 1% PS. U2OS KO cell lines expressing pINDUCER20 empty vector/UNG2-HA were generated by infecting the cells with lentivirus expressing UNG2-HA under a doxycycline-inducible promoter (pINDUCER20) and selected with G418 (400 µg/mL). Patient-derived cell lines MGH 707.1 (Raoof et al., 2019), MGH 134.1 (Crystal et al., 2014; Hata et al., 2016) and MGH143-3B (Kodack et al., 2017) were established at Massachusetts General Hospital from non-small cell lung cancer core biopsy or pleural effusion samples. All patients signed informed consent to participate in a Dana Farber/Harvard Cancer Center Institutional Review Board-approved protocol giving permission for research to be performed on their samples. Cell lines were sequenced to confirm the presence of oncogenic driver mutations.

## METHOD DETAILS

### Cloning

The human *UNG* ORF clone was purchased from Origene (Catalog#: RC222868). From this plasmid, *UNG2* was amplified and cloned into the pENTR-TOPO Gateway vector. For lentiviral expression, pENTR-TOPO-UNG2 was inserted into pINDUCER20 carrying an HA tag at the C terminus. The endogenous stop codon of *UNG2* was removed to allow in-frame expression of the C-terminal HA tag. The primers used are as follows:

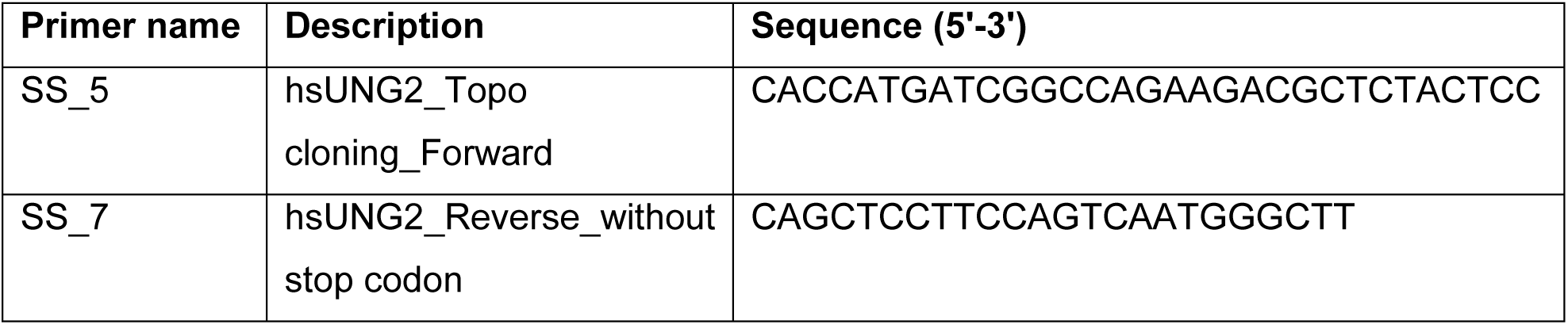

### CRISPR-Cas9 KO cell lines

UNG2 was knocked out in U2OS cells using sgRNAs previously described (Sarno et al., 2019).

Forward: caccgCGTCTTCTGGCCGATCATCC

Reverse: aaacGGATGATCGGCCAGAAGACGc

The sgRNAs targeting exon 1A of human UNG2 were cloned into PX458 vector. U2OS cells were transfected with gRNA-expressing plasmids. After 3 days, GFP-positive cells were sorted into 96-well plates as single cells by FACS and grown for 3 weeks before the cells were transferred to 24-well plates. Loss of UNG2 protein was verified by western blot using UNG antibody and 2 clones were selected for further studies.

### Plasmid transfection

For viral production and plasmid transfection, HEK293T and U2OS cells were transfected using Lipofectamine 3000 (Thermo Fisher Scientific) following manufacturer’s instructions.

### RNA interference

Reverse siRNA transfections were done using 5 nM Silencer Select pre-designed siRNAs (Thermo Fisher Scientific) with Lipofectamine RNAiMAX (Thermo Fisher Scientific) for 48 hours. We used siDESIGN Center (Horizon Discovery) to design siRNA (siUNG#4) targeting the 3′UTR region of UNG mRNA (Thermo Fisher Scientific).

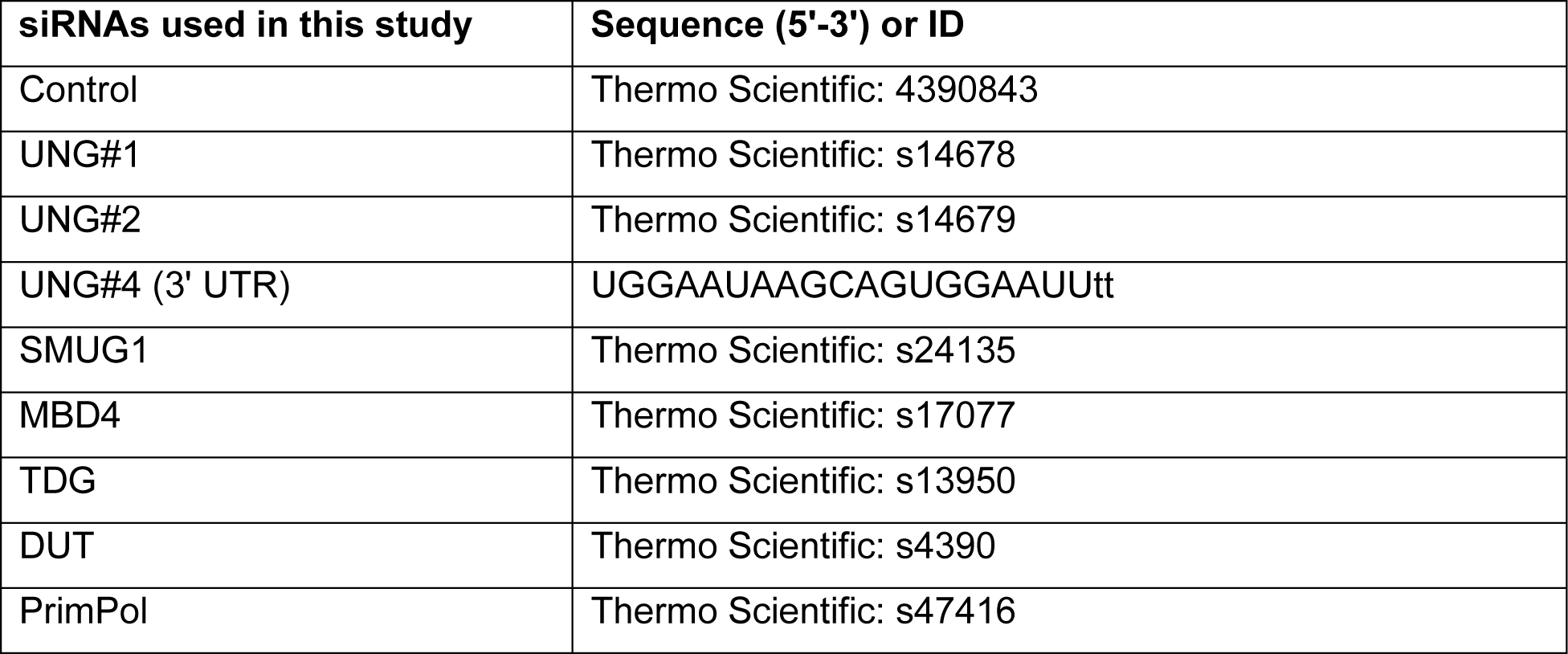

### Quantitative real-time PCR

Total RNA was isolated using NucleoSpin RNA kit for RNA purification (Macherey-Nagel; 740955). Reverse transcription was carried out with 1 μg of total RNA using random hexamer primers and the SuperScript IV reverse transcriptase kit (Thermo Fisher Scientific). Quantitative PCR was performed using Kapa SYBR Green on a Lightcycler 480 (Roche). mRNA expression relative to *GAPDH* mRNA levels was calculated using the delta-delta threshold cycle (ΔΔCT) method. The primers used are as follows:

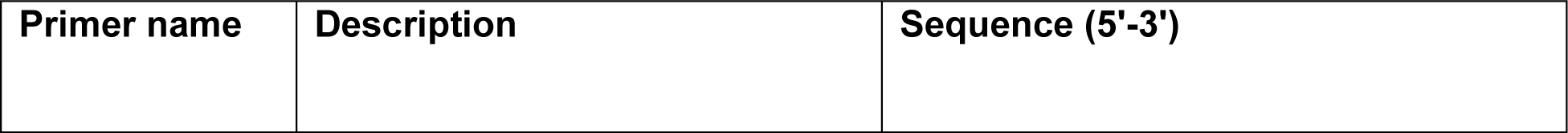
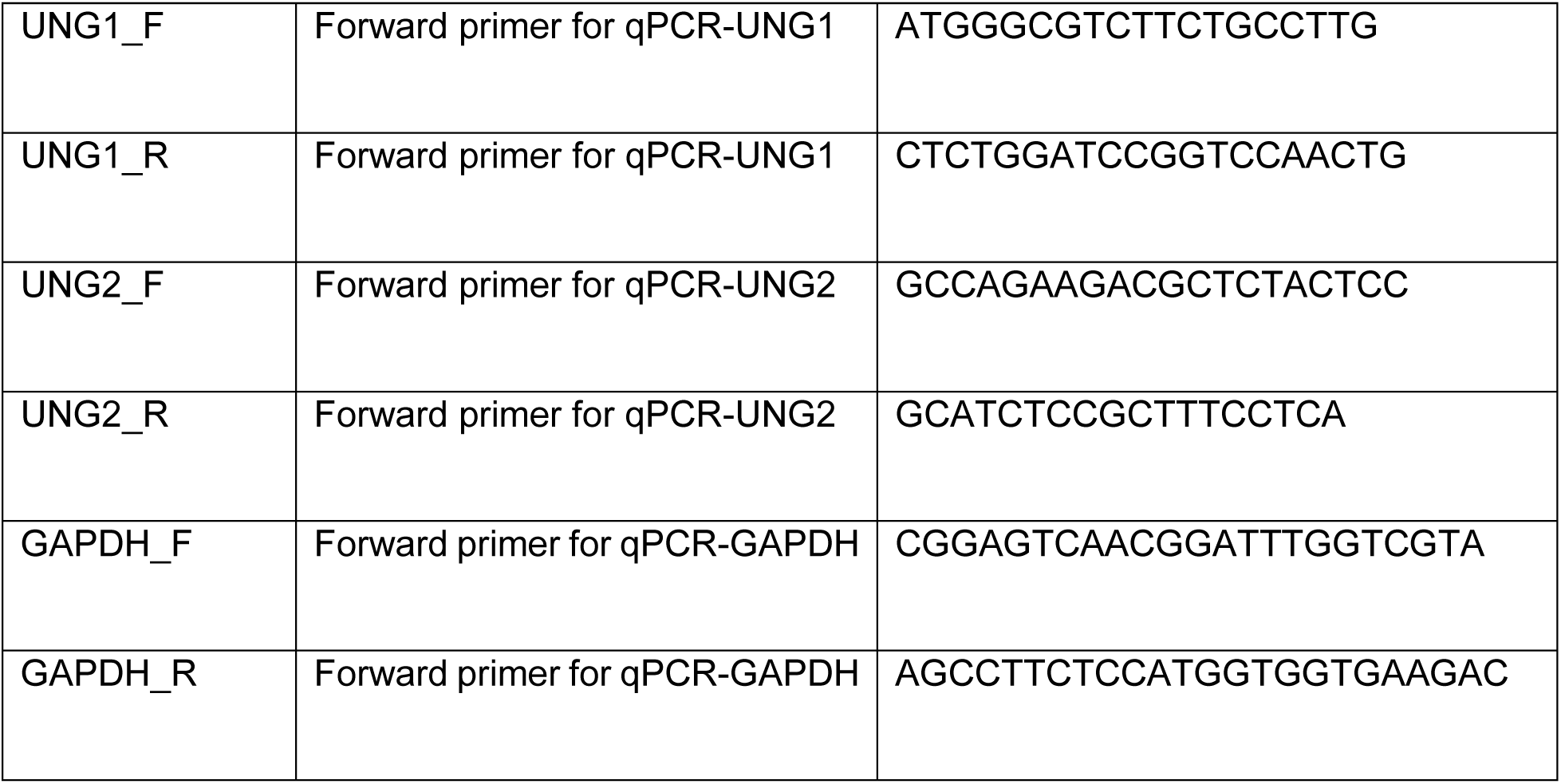

### Measurement of AP sites

AP-sites were tested in U2OS cells 48 h after transfection with control and UNG siRNA by a DNA damage assay kit (ab211154, Abcam) according to the manufacturer’s instructions.

### U-comet assay

The comet assay was performed as previously described (Paone et al., 2014). U2OS, H1299, H1838, MGH 134.1, MGH 707.1 and MGH 143-3B cells were treated with drugs as indicated, harvested, washed with PBS, and then mixed with low-melt agarose (Sigma Aldrich) at 37 °C and embedded on 1% agarose-coated Superfrost slides (Bio-Techne R&D Systems). Subsequent steps were carried out away from light. Slides were submerged in lysis buffer (2.5 M NaCl, 100 mM EDTA, 10 mM Tris, 10% DMSO, 1% Triton X-100, 8 mg/ml NaOH to pH 10) for 2 hours at 4°C, washed 3 times with enzyme buffer (20 mM Tris-HCl, 1 mM DTT, 1 mM EDTA pH 8), then treated with uracil-DNA glycosylase (Roche, UNG 0.01 unit/slide diluted in enzyme buffer) or enzyme buffer only for 1 hour at 37°C. Slides were denatured in electrophoresis buffer (1 mM EDTA, 300 mM NaOH) for 30 min at room temperature (RT), then subjected to electrophoresis at 22 V and 300 mA for 60 min at 4°C, followed by incubation in neutralization buffer (400 mM Tris, HCl to pH 7.5) for 45 min at RT. For image acquisition, slides were stained with SYBR® Gold Nucleic Acid Gel Stain (ThermoFisher) and visualized at 488 nm and 10x magnification using a Nikon 90i microscope. Comet tail moment for 100-200 cells per condition was measured using ImageJ OpenComet plugin.

### Genomic 2’-deoxyuridine measurements by High Performance Liquid Chromatography-Mass Spectrometry (LC-MS)

Genomic DNA was purified from 2-3 million cells using Qiagen DNeasy Blood & Tissue Kit (Product #69504), incorporating Proteinase K and RNase A treatment per manufacturer’s recommendations. DNA was eluted in DNase/RNase-free water and dried over gaseous N_2_. Total DNA yields were measured using Invitrogen Qubit dsDNA BR assay (Product #Q32851) and recorded for subsequent normalization of mass spectrometry data. Typical yields ranged from 3-7 µg of DNA. dU content was measured by the treatment of DNA with Uracil DNA Glycosylase (UDG), which generates free uracil nucleobase. DNA was resuspended in UDG reaction buffer (10 mM Tris HCl, pH 8.0@25°C, 1 mM EDTA, 1 mM DTT) containing 50 fmol of stable, isotopically-labeled 1,3-15N2 Uracil (Cambridge Isotope Laboratories, Catalog number NLM-637-0; Lot Number PR32662B). *E. Coli* UDG (New England Biolabs Inc., Catalog number M0280L, Lot number 10151474) was dialyzed twice in 1000 sample volumes of UDG reaction buffer at 4°C for 12 hours to remove glycerol from enzyme concentration. 10 U of UDG enzyme was added to 100 µl of genomic DNA resuspended in UDG reaction buffer and incubated at 37°C for 2 hours. Samples were then spin-filtered with Amicon Ultra 0.5 ml Centrifugal filters with 3 kilodalton molecular weight cutoff (Catalog number UFC500396). Eluate was dried over gaseous N_2_ and stored at -80°C prior to analysis by High Performance Liquid Chromatography-Mass Spectrometry (LC-MS).

On the day of LC-MS analysis, samples were resuspended in 20 µl HPLC-grade Water (Sigma-Aldrich, Catalog number 270733-4L). Metabolites were measured using a Dionex UltiMate 3000 ultrahigh-performance liquid chromatography system connected to a Q Exactive benchtop Orbitrap mass spectrometer, equipped with an Ion Max source and a HESI II probe (Thermo Fisher Scientific). Samples were separated by chromatography by injecting 2 μl of sample on a SeQuant ZIC-pHILIC Polymeric column (2.1 × 150 mm 5 μM, EMD Millipore). The flow rate was set to 150 μl/min, temperatures were set to 25 °C for column compartment and 4°C for autosampler sample tray. Mobile Phase A consisted of 20 mM ammonium carbonate, 0.1% ammonium hydroxide. Mobile Phase B was 100% acetonitrile. The mobile phase gradient (%B) was set in the following protocol: 0–20 min, linear gradient from 80% to 20% B; 20–20.5 min, linear gradient from 20% to 80% B; 20.5–28 min, hold at 80% B. Mobile phase was introduced into the ionization source set to the following parameters: sheath gas, 40; auxiliary gas, 15; sweep gas, 1; spray voltage, −3.1 kV; capillary temperature, 275 °C; S-lens RF level, 40; probe temperature, 350 °C. Metabolites were monitored with a full-scan and additional narrow range scan (110.5–113.5 m/z) in negative mode only. The resolution was set at 70,000, the AGC target at 1,000,000 and the maximum injection time at 20 ms. Relative quantitation of metabolites was performed with XCalibur QuanBrowser 2.2 (Thermo Fisher Scientific) using a 5 ppm mass tolerance and referencing a retention time for uracil from an in-house library of chemical standards. Total ion counts were normalized to the internal 1,3-15N2-labelled uracil standard and nanograms of input genomic DNA per sample.

### DNA fiber analysis

Exponentially growing U2OS, H1299 or H1838 cells were first pulse labeled with 100 μM CldU, washed twice with equilibrated complete media and then labeled with 250 μM IdU under the conditions specified in the figure legends. Collected cells were resuspended in cold PBS (1×10^6^ cell per milliliter), and 3 μL of cell suspension was mixed with spreading buffer (8 μL) (0.5% SDS, 200 mM Tris-HCl pH 7.4, 50 mM EDTA) and let spread by gravity on a tilted glass slide. DNA fibers were fixed in methanol:acetic acid (3:1) for 10 min, denatured in 2.5 N HCl for 60 min, and blocked in 3% BSA/0.05% Tween-20 for 30 min at 37°C. CldU and IdU detection were done using rat anti-BrdU (1:100; Abcam) and mouse anti-BrdU (1:50; BD Biosciences) for 1 hour at 37°C followed by Alexa 488 anti-mouse (1:100; Jackson ImmunoResearch) and Cy3 anti-rat (1:100; Jackson ImmunoResearch) for 30 minutes at 37°C. Slides were mounted with Prolong Gold and dried overnight. Fibers were imaged with a 60X objective on a Nikon 90i microscope and quantified using FIJI.

S1 nuclease experiments were essentially performed as described previously (Quinet et al., 2017). After IdU labeling, cells were permeabilized with CSK100 buffer (100 mM NaCl, 10 mM MOPS pH7, 3 mM MgCl2, 300 mM sucrose, 0.5% Triton X-100) for 10 minutes at room temperature (RT) and then washed with 1x PBS carefully. The cells were washed once with S1 nuclease buffer pH4.6 (30 mM NaAc, 10 mM ZnAc, 5% glycerol, 50 mM NaCl) and subsequently incubated with S1 buffer containing S1 nuclease (20 U/ mL) for 30 minutes at 37°C. The cells were washed once with 1x PBS containing 0.1% BSA to help precipitate nuclei, then collected and processed for staining as described previously.

### Uracil excision activity assay

U2OS cells were transfected with control or UNG siRNA and lysed in native lysis buffer (25 mM HEPES pH 7.9, 10% glycerol, 150 mM NaCl, 0.5% Triton X-100, 1 mM EDTA, 1 mM MgCl2, 0.2 µg/ml RNase A and protease inhibitors) after 48 h. Cell lysates were sonicated, incubated for 30 min at 4°C, and then centrifuged for 10 min at 13,000 RPM at 4°C. Protein concentration was determined by BCA Assay (Thermo Fisher Scientific) and extracts were normalized. 20 uL aliquots were prepared and flash frozen in liquid nitrogen and stored in -80C. A fresh vial of extract was used for each experiment.

For the activity assay, normalized amounts of cell extracts were incubated with 0.4 µM DNA hairpin substrate for 1 h at 37°C in reaction buffer (50 mM Tris pH 7.5, 0.1 mg/ml RNase A, and 5 mM EDTA). DNA oligonucleotide probe was synthesized by ThermoFisher Scientific having the following sequence: 5’FAM-GCAAGCCTTUGGCTTGCTGA. For the positive control, DNA substrate was incubated with 1 unit of purified Uracil-DNA glycosylase (New England BioLabs) instead of cell extracts in the same reaction buffer. After 1 h, 100 mM NaOH was added to the reaction and further incubated at 95°C for 30 min to cleave AP-sites. Gel loading buffer (0.5% Orange G in Formamide) at 1:1 concentration was then added to the reaction and further incubated at 95°C for 10 min, spun down and kept on ice for 5 min. A 20% denaturing polyacrylamide gel (8 M urea, 1× TAE buffer) was pre-run in 1X TAE buffer for 10 min and 10 uL samples were loaded. DNA cleavage was monitored by running the gel for 90 min at 150 V. The gel was analysed on a Chemidoc imaging system (BioRad) with ImageLab v6.0.1 software. Quantification was performed using Fiji.

### Import of dNTPs into live cells

U2OS cells were treated with 2 µM Bio-tracker (BT) and indicated concentrations of Cy3-dUTP, dUTP or dTTP for 24 h in Leibovitz’s L-15 Medium supplemented with 10% FBS and 1% PS. For cell survival assays, after 24 h treatment with BT and Cy3-dUTP, the cells were seeded in 96-well plates in fresh DMEM supplemented with L-glutamine, 10% FBS, and 1% PS and indicated concentrations of ATRi and cultured for 5-7 days.

### Immunofluorescence

To monitor DNA synthesis, U2OS cells were pulse-labeled with 10 μM EdU for 30 minutes and then treated as described in the figure legends. Cells were pre-extracted with 1x PBS containing 0.2% Triton X-100 for 2 minutes prior to fixation with 3% paraformaldehyde/2% sucrose for 10 min at RT. Following two washes with 3% BSA in 1x PBS, coverslips were incubated with 2 mM CuSO4, 10 mM sodium ascorbate, and 1 μM picolyl-azide AF647 in 1x TBS for 30 minutes at RT, followed by two washes with 3% BSA in 1x PBS. Coverslips were then incubated with the IF blocking buffer (1% BSA+0.5% T-X100 in 1x PBS) for 60 min at RT. Primary antibodies diluted in blocking buffer were added to the cells, and incubation continued for 2 hours at RT. After two washes with PBST (1x PBS with 0.5% T-X100), cells were incubated in the dark with secondary antibodies, diluted to 1:250 in blocking buffer, for 1 hour at RT. Finally, after two washes with PBST, cells were stained with DAPI for 5 minutes and mounted on slides with Prolong Gold. Images were captured with a Nikon 90i microscope and quantified using FIJI.

For 8-oxoG immunostaining, coverslips were fixed in 3% paraformaldehyde/2% sucrose for 20 min at RT, followed by permeabilization in 1x PBS containing 0.5% Triton X-100 for 5 minutes. Next, coverslips were treated with RNAse buffer (1 mM EDTA, 10 mM Tris-HCl (pH 7.5), 0.4 mM NaCl, and 100 µg/mL RNAse (Invitrogen) for 1 hour at 37°C, followed by denaturation in 2.5 N HCl for 30 min at RT. After two washes and 10 min incubation in neutralization buffer (50 mM Tris–HCl pH 8.8) at RT, coverslips were blocked with 4% BSA+0.1% T-X100 in PBS for 1 hour at RT. Primary antibodies diluted in blocking buffer were added to the cells, and incubation continued overnight at 4°C. After two washes with PBST (1x PBS with 0.5% T-X100), cells were incubated in the dark with secondary antibodies, diluted to 1:250 in blocking buffer, for 1 hour at RT. Finally, after two washes with PBST, cells were stained with DAPI for 5 minutes and mounted on slides with Prolong Gold. Images were captured with a Nikon 90i microscope and quantified using FIJI.

### Proximity ligation assay (PLA)

PLA was performed as described previously (Matos et al., 2020). Briefly, cells were treated with CSK extraction buffer (0.2% Triton X-100, 20 mM HEPES-KOH pH 7.9, 100 mM NaCl, 3 mM MgCl2, 300 mM sucrose, 1 mM EGTA), fixed with PFA and methanol, and then permeabilized with 1x PBS containing 0.5% Triton-x100. The cells were treated with RNaseA (10 ug/ml) for 1 hour at 37 °C and then washed and blocked with 3% BSA in PBST buffer for 1 hour. The cells were incubated with the primary antibodies diluted at 1:500 at 4°C overnight. After three washes with 1x PBST, the cells were incubated with anti-mouse minus and anti-rabbit plus PLA probes (PLA kit from Sigma) at 37°C for 1 hour. The cells were washed with PLA buffer A and incubated with ligation buffer containing ligase (PLA kit) for 30 minutes at 37°C, then washed and incubated with amplification buffer with polymerase (PLA kit) at 37°C for 1 hour. The cells were washed twice with PLA buffer B (PLA kit) and then three times with PBST buffer containing DAPI. Images were captured with a Nikon 90i microscope.

### Immunoblots

Cells were harvested and lysed in RIPA buffer supplemented with 1x protease inhibitors. Protein concentrations were normalized using Bradford assay and mixed 1:1 with 2× SDS PAGE loading buffer (100 mM Tris at pH 6.8, 12% glycerol, 3.5% SDS, 0.2 M DTT). Samples were boiled for 10 minutes, loaded on Bolt Bis-Tris Plus 4%–12% gels, and run at 100 V for 90 min. Proteins were transferred onto PVDF membranes using a CBS Scientific electrophoretic blotting liquid transfer system (EBX-700) for 1 hour at 300 mA. Membranes were then blocked in Tris-buffered saline with 0.05% Tween-20 (TBS-T) and 3% BSA for 60 min at room temperature. Membranes were then immunoblotted with primary antibodies, diluted in blocking buffer, overnight at 4°C with mild shaking. The PrimPol antibody was kindly provided by the Méndez laboratory. Membranes were washed twice for 5 min with TBST and incubated for 2-3 hours at 4°C with secondary antibodies conjugated to horseradish peroxidase. Membranes were washed twice for 5 minutes with TBST and an enhanced chemiluminescence (ECL Bio-Rad 1705061) solution was applied. Signals were detected using a Chemidoc imaging system (Bio-Rad) with ImageLab v6.0.1. software.

### Chromatin fractionation

Cells were harvested and washed with PBS. Cytosolic proteins were removed by incubation of cells in hypotonic buffer (10 mM HEPES-KOH pH 7.5, 5 mM KCl, 1.5 mM MgCl2, 1 mM DTT and 0.5% NP-40) for 15 minutes on ice and centrifuged at 13000 rpm for 5 minutes. The nucleus-enriched pellets were resuspended in fraction buffer (10 mM HEPES-KOH pH 7.5, 5 mM KCl, 1.5 mM MgCl2, 1 mM DTT and 0.5% NP-40 and 0.5 M NaCl), incubated on ice for 10 minutes and centrifuged at 13000 rpm for 5 minutes. The supernatants containing soluble nuclear proteins was discarded and the chromatin-enriched pellets were washed once with fraction buffer. The final pellets enriched for stably chromatin-bound proteins were incubated in 2x Laemmle sample buffer for 10 minutes at 90°C and used for western blotting.

### Cell viability assay

Cell viability assays were performed in 96-well format and cells were treated as described in the figure legends. To determine the number of viable cells, CellTiter-Glo was used according to manufacturer’s instructions.

### *In vivo* drug response

2 x 10^6^ H1299 non-small cell lung carcinoma cells were injected into the right flank of female NOD scid gamma (NSG) mice (Jackson Laboratory) in equal proportions with Matrigel Basement Membrane Matrix (Corning, CLS354234). Tumors were allowed to reach 100±5 mm^3^ prior to beginning treatment. Four mice were treated per cohort. Mice were administered either intraperitoneal injections of Pemetrexed (Selleckchem, S5971) in 0.9% NaCl (75 mg/kg), AZD6738 (Selleckchem, S7693) in 5% DMSO and 40% propylene glycol (50 mg/kg), or both. Control mice were administered equal amounts of DMSO via the same route. Mice were treated every other day for a total of 15 treatments. Tumors were measured externally using calipers and tumor volume was calculated using the following formula: (length x width^2^)/2. Mice were weighed once a week to monitor changes in weight.

## QUANTIFICATION AND STATISTICAL ANALYSIS

FIJI software was used to quantify immunofluorescence intensities and measure IdU tract length. Statistical analyses were performed using the GraphPad Prism software. Statistical parameters are shown in figures and described in figure legends. For DNA fiber analysis, p value significance was defined as follows: (ns) non-significant; (*) P ≤ 0.05; (**) P ≤ 0.01; (***) P ≤ 0.001.

