## Supplementary Figures and Legends for "Unprocessed Genomic Uracil as a Source of DNA Replication Stress in Cancer Cells"

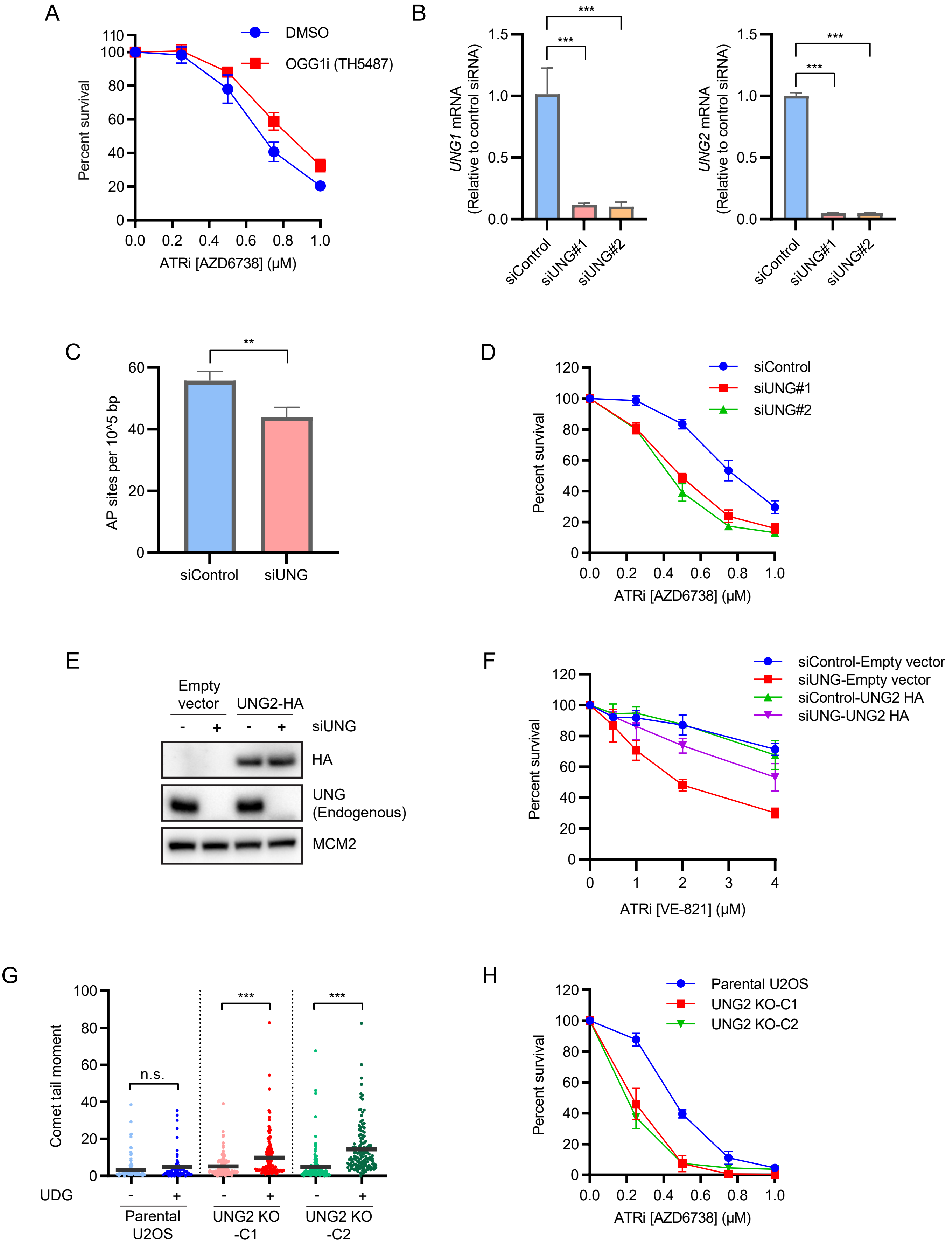


Supplementary Figure 1

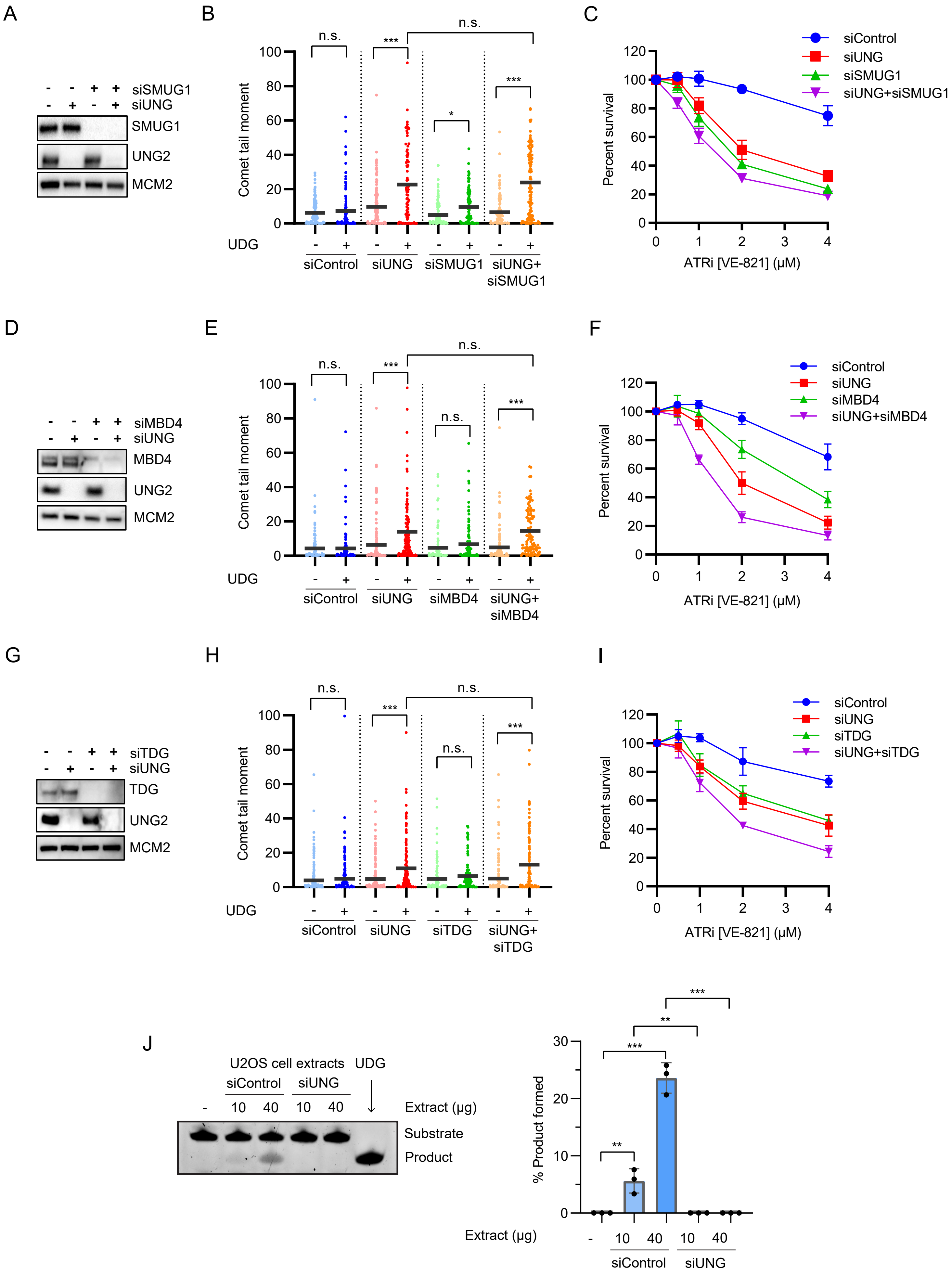


Supplementary Figure 2

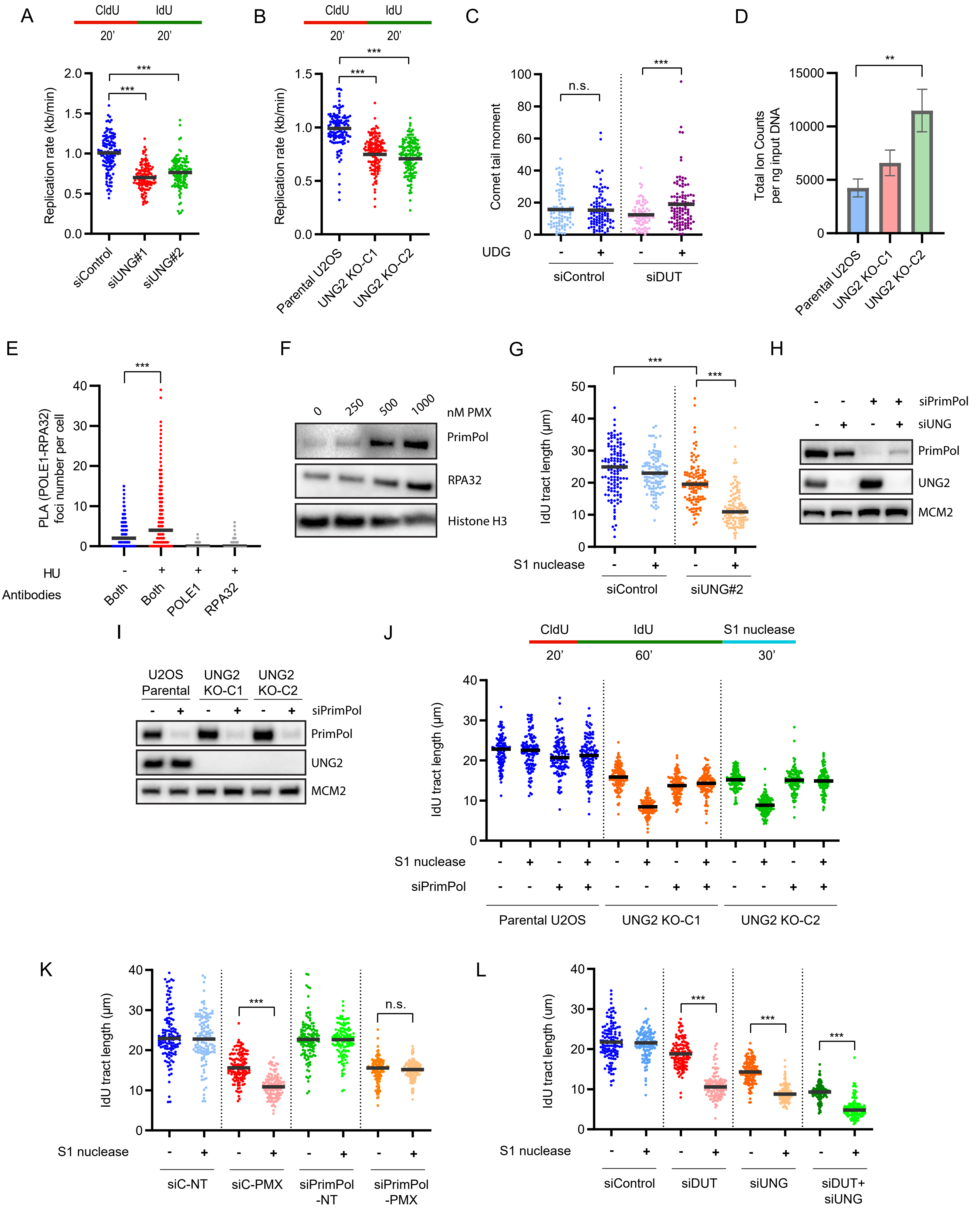


Supplementary Figure 3

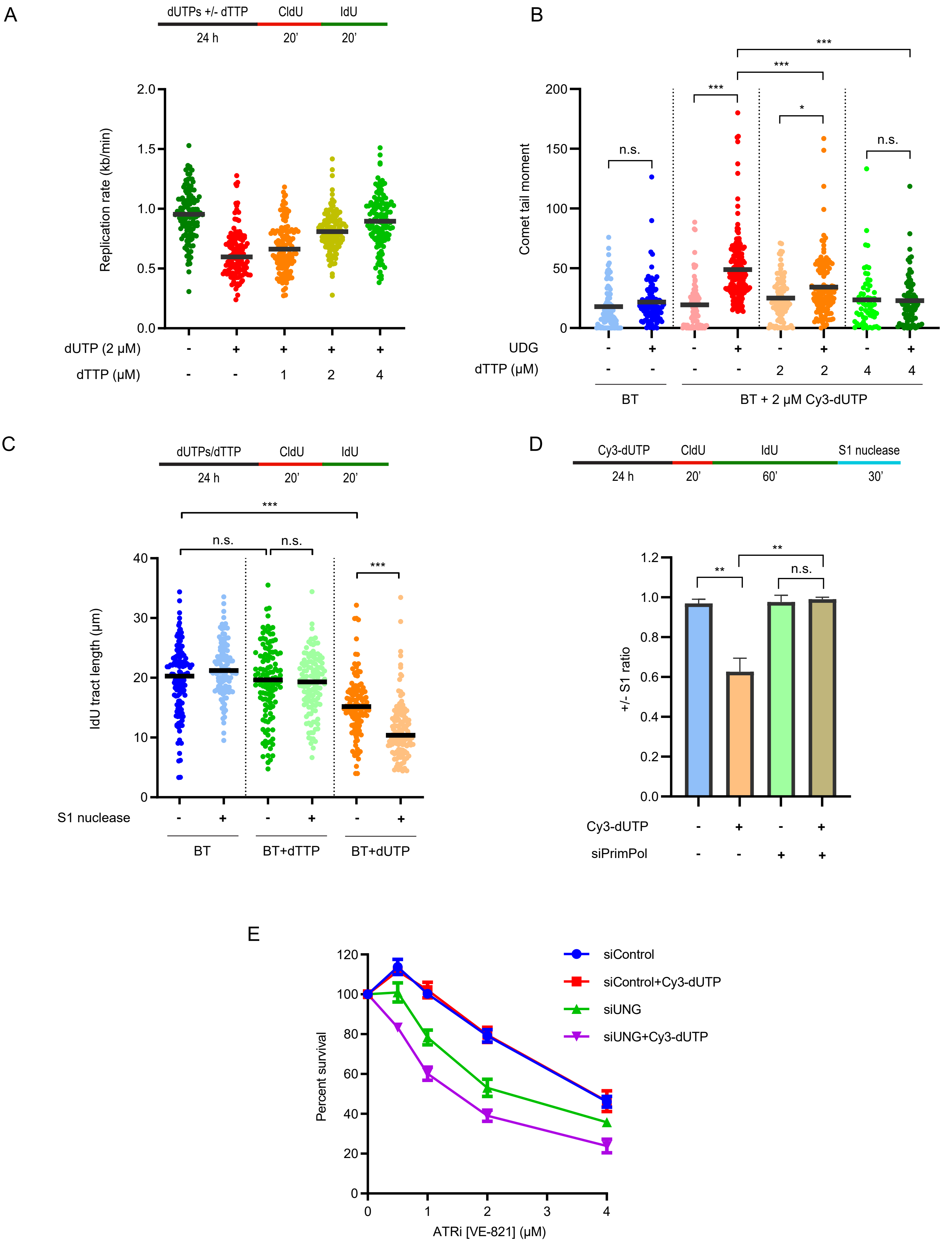


Supplementary Figure 4

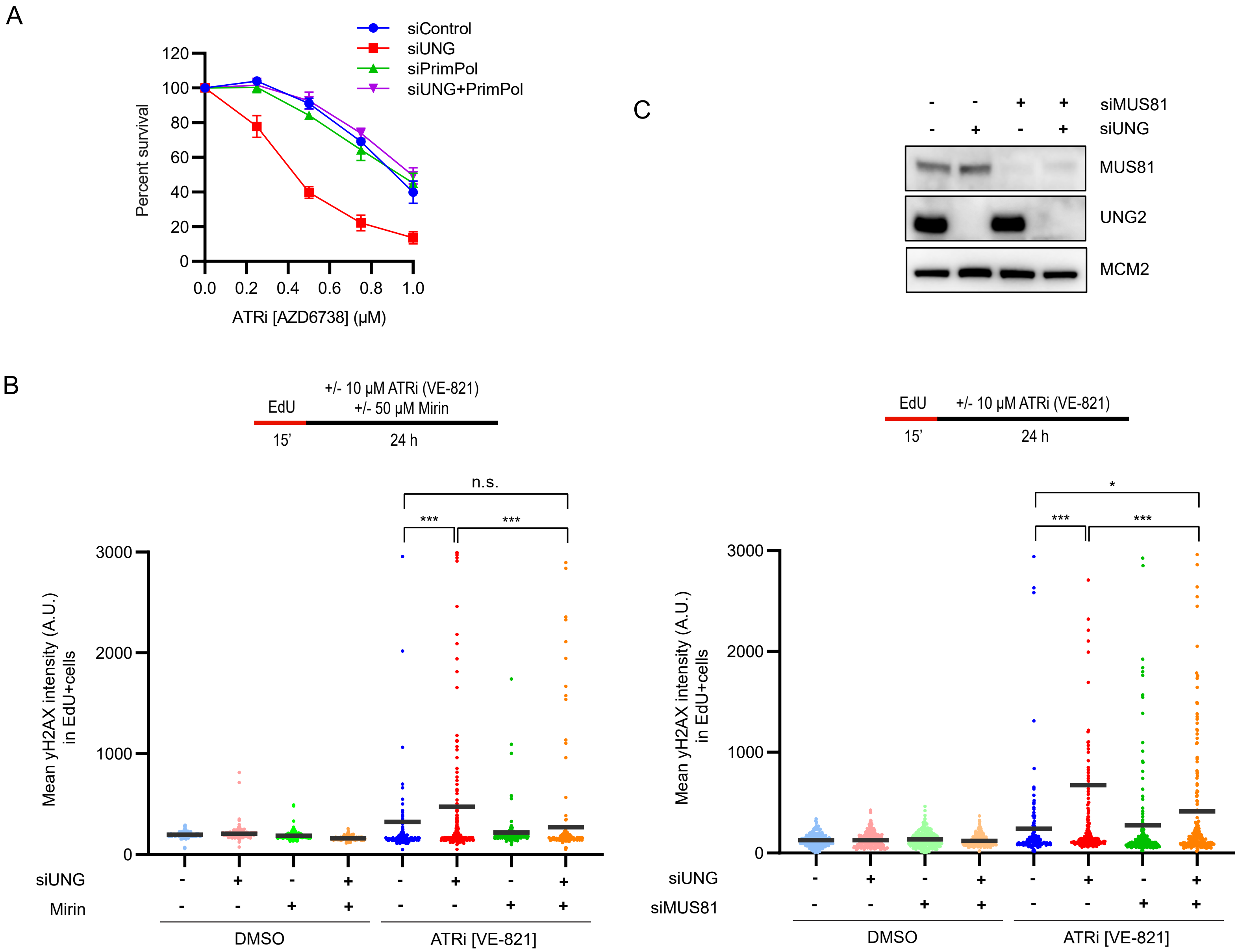


Supplementary Figure 5

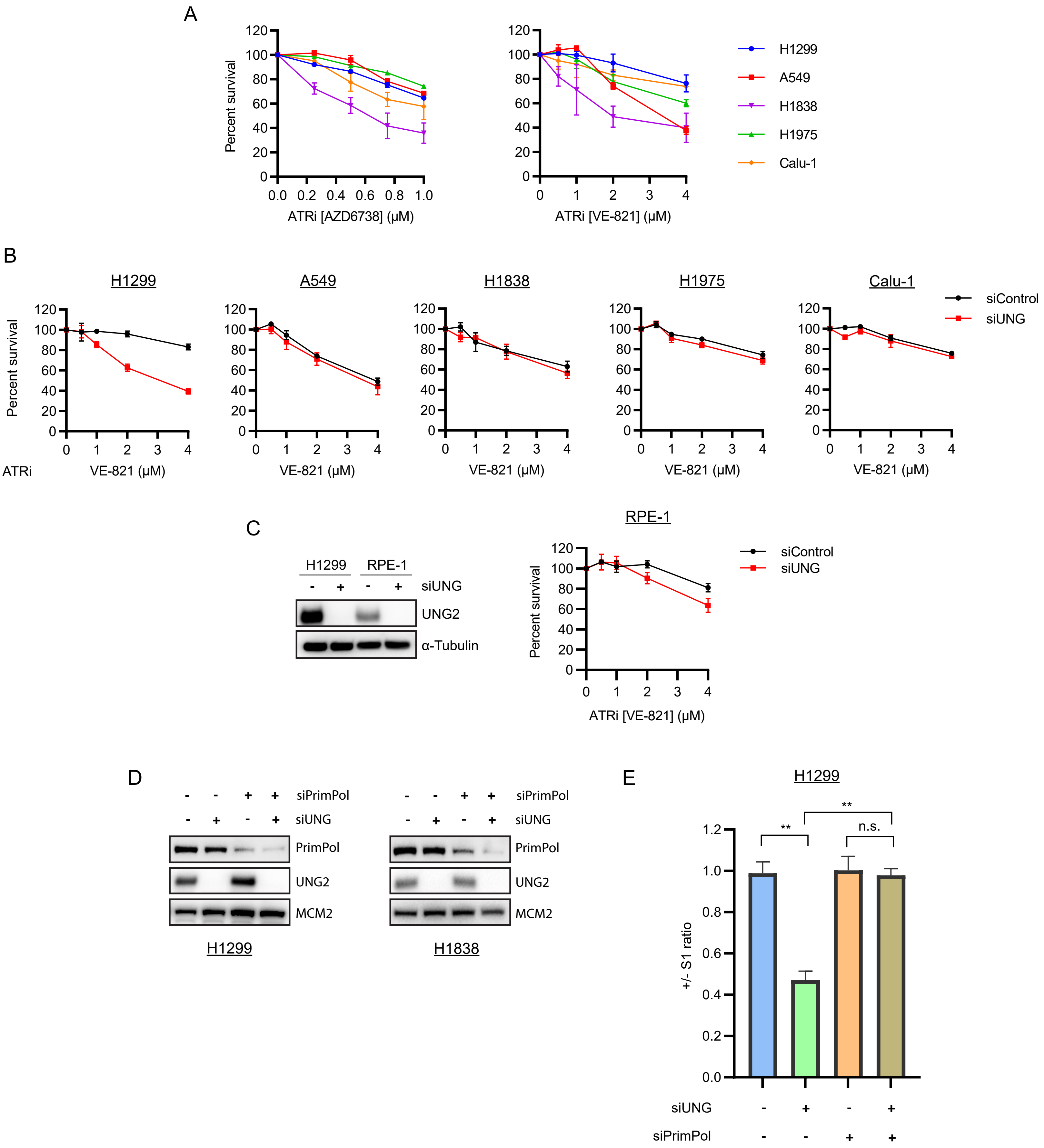


Supplementary Figure 6

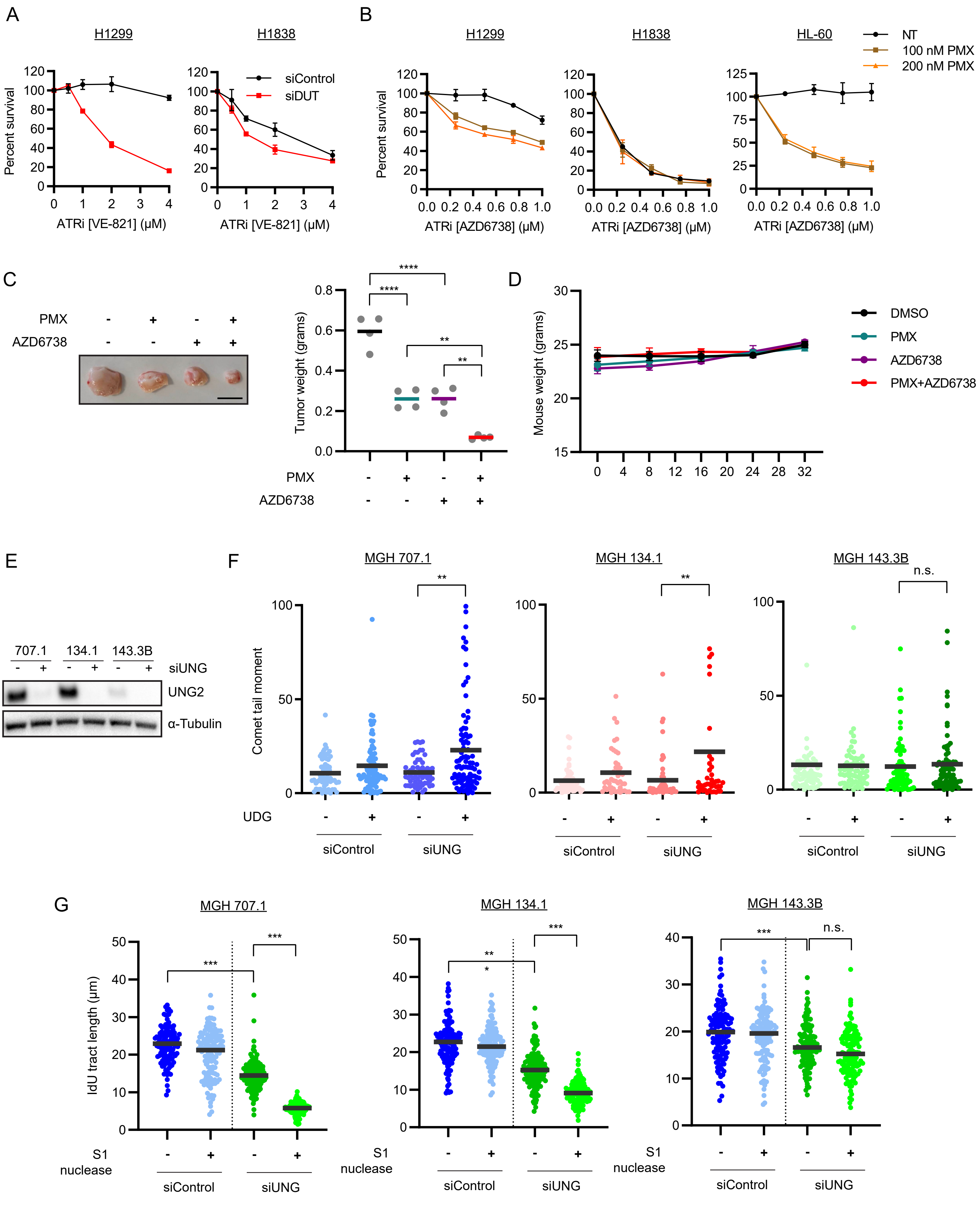


Supplementary Figure 7

**Supplementary Figure legends**

**Supplementary Figure 1 (associated with Fig. 1). Processing of genomic uracil by uracil glycosylase UNG2 reduces ATRi sensitivity.**

(A) U2OS cells were treated with DMSO or 10 µM TH5487 and indicated concentrations of ATRi (AZD6738) for 5-7 days. Cell viability was determined using CellTiter-Glo and normalized to the cells untreated with ATRi under each condition. Data are shown as mean ± s.d. (n=2 independent experiments).

(B) Fold change in *UNG1* and *UNG2* mRNA levels in U2OS cells transfected with indicated siRNAs relative to control siRNA-treated cells. *UNG1* and *UNG2* mRNA levels were normalized to *GAPDH* mRNA levels. Data are shown as mean ± s.d. (n=3 independent experiments).

(C) U2OS cells were transfected with control or UNG#1 siRNA for 48 h and AP-sites were measured using Aldehyde Reactive Probe (ARP). Data are shown as mean ± s.d. (n=3 independent experiments).

(D) U2OS cells were transfected with indicated siRNAs and treated with indicated concentrations of ATRi (AZD6738) for 5-7 days. Cell viability was determined using CellTiter-Glo and normalized to the cells untreated with ATRi under each condition. Data are shown as mean ± s.d. (n=2 independent experiments).

(E) U2OS cells that inducibly express UNG2-HA were treated with 400 ng/ml doxycycline (DOX) to induce the expression of UNG2-HA. After 24 h, cells were transfected with UNG 3' UTR specific siRNA. At 48 h post-transfection, levels of the indicated proteins were analyzed by western blot. Note that UNG antibody does not detect UNG2-HA. MCM2 is shown as the loading control in western blot.

(F) At 48 h post-siRNA transfection, the cells described in (C) were treated with indicated concentrations of ATRi (AZD6738) and treatments continued for 5-7 days. Cell viability was determined using CellTiter-Glo and normalized to the cells untreated with ATRi under each condition. Data are shown as mean ± s.d. (n=2 independent experiments).

(G) Wild-type (WT) or *UNG2* KO U2OS cells were analyzed by U-comet assay. Dot plots represent comet tail moment in individual cells, and bars display the mean in cell populations (n>50 cells per condition). One of two independent experiments is shown.

(H) WT and *UNG2* KO U2OS cells were treated with indicated concentrations of ATRi (AZD6738) for 5-7 days. Cell viability was determined using CellTiter-Glo and normalized to the cells untreated with ATRi under each condition. Data are shown as mean ± s.d. (n=2 independent experiments).

**Supplementary Figure 2 (associated with Fig. 1). Genomic uracil is not efficiently processed in cells lacking UNG2.**

(A) U2OS cells were transfected with UNG#1 and SMUG1 siRNA. At 48 h post-transfection, levels of the indicated proteins were analyzed by western blot. MCM2 is shown as the loading control in western blot.

(B) U2OS cells transfected with UNG#1 and SMUG1 siRNA were analyzed by U-comet assay. Dot plots represent comet tail moment in individual cells, and bars display the mean in cell populations (n>100 cells per condition). One of two independent experiments is shown.

(C) U2OS cells were transfected with UNG#1 and SMUG1 siRNA and treated with indicated concentrations of ATRi (VE-821) for 5-7 days. Cell viability was determined using CellTiter-Glo and normalized to the cells untreated with ATRi under each condition. Data are shown as mean ± s.d. (n=3 independent experiments).

(D) U2OS cells were transfected with UNG#1 and MBD4 siRNA. At 48 h post-transfection, levels of the indicated proteins were analyzed by western blot. MCM2 is shown as the loading control in western blot.

(E) U2OS cells transfected with UNG#1 and MBD4 siRNA were analyzed by U-comet assay. Dot plots represent comet tail moment in individual cells, and bars display the mean in cell populations (n>100 cells per condition). One of two independent experiments is shown.

(F) U2OS cells were transfected with UNG#1 and MBD4 siRNA and treated with indicated concentrations of ATRi (VE-821) for 5-7 days. Cell viability was determined using CellTiter-Glo and normalized to the cells untreated with ATRi under each condition. Data are shown as mean ± s.d. (n=3 independent experiments).

(G) U2OS cells were transfected with UNG#1 and TDG siRNA. At 48 h post-transfection, levels of the indicated proteins were analyzed by western blot. MCM2 is shown as the loading control in western blot.

(H) U2OS cells transfected with UNG#1 and TDG siRNA were analyzed by U-comet assay. Dot plots represent comet tail moment in individual cells, and bars display the mean in cell populations (n>100 cells per condition). One of two independent experiments is shown.

(I) U2OS cells were transfected with UNG#1 and TDG siRNA and treated with indicated concentrations of ATRi (VE-821) for 5-7 days. Cell viability was determined using CellTiter-Glo and normalized to the cells untreated with ATRi under each condition. Data are shown as mean ± s.d. (n=3 independent experiments).

(J) U2OS cells were transfected with control or UNG#1 siRNA and cell extracts were used to measure uracil excision activity on uracil-containing DNA hairpin substrate. Left, Representative gel image showing the comparison of uracil excision activity in cell extracts prepared from control and UNG2 knockdown cells. UNG2 depleted cells show no detectable uracil excision activity. Right, Quantification of band intensities of reaction products from the gel. Data are shown as mean ± s.d. (n=3 independent experiments).

**Supplementary Figure 3 (associated with Fig. 2 and Fig. 3). Unprocessed genomic uracil impedes replication forks and induces PrimPol-generated ssDNA gaps.**

(A) U2OS cells were transfected with the indicated siRNAs for 48 h, and replication rate was analyzed by DNA fiber assay as indicated (n=125 fibers per condition). One of two independent experiments is shown.

(B) Replication rate in WT or *UNG2* KO U2OS cells was analyzed by DNA fiber assay as indicated (n=125 fibers per condition). One of two independent experiments is shown.

(C) U2OS cells transfected with UNG#1 and DUT siRNA were analyzed by U-comet assay. Dot plots represent comet tail moment in individual cells, and bars display the mean in cell populations (n>50 cells per condition). One of two independent experiments is shown.

(D) Parental and *UNG2* KO U2OS cells were treated with 200 nM Pemetrexed (PMX) for 24 h and genomic uracil incorporation was measured by LC-MS. Data are shown as mean ± s.d. (n=3).

(E) U2OS cells were treated with or without 2 mM hydroxyurea (HU) for 1 h and then analyzed by PLA using anti-POLE1 and anti-RPA32 antibodies, either alone or in combination. Quantification of PLA foci in individual cells is shown (n>1200 cells per condition).

(F) U2OS cells were treated with indicated concentrations of PMX for 24 h. Chromatin fractions were prepared and blotted for PrimPol and RPA32. Histone H3 is shown as the loading control.

(G) U2OS cells were transfected with control or UNG#2 siRNA for 48h, and then analyzed by DNA fiber assay as in Fig. 3D. DNA fibers were treated with S1 nuclease (20 U/mL) for 30 min at 37°C, and the length of IdU replication tracts (n=125 fibers per condition) was measured. One of two independent experiments is shown.

(H) U2OS cells were transfected with UNG#1 and PrimPol siRNA for 48 h. Levels of PrimPol and UNG2 in whole-cell extracts were analyzed by western blot. MCM2 is shown as the loading control.

(I) WT or *UNG2 KO* U2OS cells were transfected with control or PrimPol siRNA for 48 h. Levels of PrimPol and UNG2 in whole-cell extracts were analyzed by western blot. MCM2 is shown as the loading control.

(J) DNA synthesis in cells as in (H) was analyzed by DNA fiber assay as indicated. DNA fibers were treated with S1 nuclease (20 U/mL) for 30 min at 37°C, and the length of IdU replication tracts (n=125 fibers per condition) was measured. One of two independent experiments is shown.

(K) U2OS cells transfected with control or PrimPol siRNA were treated with 200 nM PMX for 24 h and analyzed as in (I). The length of IdU replication tracts (n=125 fibers per condition) was measured. One of two independent experiments is shown.

(L) U2OS cells transfected with UNG#1 and dUTPase (DUT) siRNA were analyzed as in (I). The length of IdU replication tracts (n=125 fibers per condition) was measured. One of two independent experiments is shown.

**Supplementary Figure 4 (associated with Fig. 4). Exogenous dUTP induces ssDNA gaps and replication stress.**

(A) U2OS cells were treated with 2 μM Biotracker (BT), 2 μM dUTP and indicated concentrations of dTTP for 24 h, and replication rate was analyzed by DNA fiber assay as indicated (n=125 fibers per condition). One of two independent experiments is shown.

(B) U2OS cells were treated with 2 μM BT, 2 μM Cy3-dUTP and indicated concentrations of dTTP for 24 h and analyzed by U-comet assay. Dot plots represent comet tail moment in individual cells, and bars display the mean in cell populations (n>65 cells per condition). One of two independent experiments is shown.

(C) U2OS cells were treated with 2 µM BT and 2 µM dUTP/dTTP for 24 h, and replication rate was analyzed by DNA fiber assay as indicated (n=125 fibers per condition). One of two independent experiments is shown.

(D) Ratio of IdU tracts from S1-treated to -untreated fibers in Fig. 4E. Data are displayed as means ± s.d. from two independent experiments.

(E) U2OS cells transfected with control or UNG#1 siRNA were treated with 2 µM BT and 2 µM Cy3-dUTP for 24 h. Cells were then allowed to recover in fresh media and treated with indicated concentrations of ATRi (VE-821) for 5-7 days. Cell viability was determined using CellTiter-Glo and normalized to the cells untreated with ATRi under each condition. Data are shown as mean ± s.d. (n=2 independent experiments).

**Supplementary Figure 5 (associated with Fig. 5). PrimPol-generated ssDNA gaps increase ATRi sensitivity in UNG-deficient cells.**

(A) U2OS cells were transfected with UNG#1 and PrimPol siRNA and treated with indicated concentrations of ATRi (AZD6738) for 5-7 days. Cell viability was determined using CellTiter-Glo and normalized to the cells untreated with ATRi under each condition. Data are shown as mean ± s.d. (n=2 independent experiments).

(B) U2OS cells were transfected with UNG#1 and pulse labeled with 5 µM EdU for 15 min, followed by treatment with 10 µM ATRi (VE-821) +/- 50 µM MRE11i (Mirin) for 24 h. γH2AX intensity in EdU-positive cells was quantified (n>250 cells per condition). One of two independent experiments is shown.

(C) U2OS cells were transfected with UNG#1 and MUS81 siRNA and pulse labeled with 5 µM EdU for 15 min, followed by treatment with 10 µM ATRi (VE-821) for 24 h. γH2AX intensity in EdU-positive cells was quantified (n>250 cells per condition). One of two independent experiments is shown.

**Supplementary Figure 6 (associated with Fig. 6). High UNG expression in cancer cells is associated with uracil-induced replication stress.**

(A) A panel of lung cancer cell lines were treated with indicated concentrations of ATRi (VE-821 or AZD6738) for 5-7 days. Cell viability was determined using CellTiter-Glo and normalized to the cells untreated with ATRi under each condition. Data are shown as mean ± s.d. (n=2 independent experiments).

(B) Indicated cell lines were transfected with control or UNG#1 siRNA and treated with indicated concentrations of ATRi (VE-821) for 5-7 days. Cell viability was determined using CellTiter-Glo and normalized to the cells untreated with ATRi under each condition. Data are shown as mean ± s.d. (n=4 independent experiments).

(C) Left, H1299 and RPE-1 cells were transfected with control or UNG#1 siRNA for 48 h. Levels of UNG2 in whole-cell extracts were analyzed by western blot. α-Tubulin is shown as the loading control. Right, RPE-1 cells transfected with control or UNG#1 siRNA were treated with indicated concentrations of ATRi (VE-821) for 5-7 days. Cell viability was determined using CellTiter-Glo and normalized to the cells untreated with ATRi under each condition. Data are shown as mean ± s.d. (n=3 independent experiments).

(D) H1299 and H1838 cells were transfected with UNG#1 and PrimPol siRNA for 48 h. Levels of PrimPol and UNG2 in whole-cell extracts were analyzed by western blot. MCM2 is shown as the loading control.

(E) DNA fibers from H1299 cells were analyzed as shown in Fig. 6C. Ratio of IdU tracts from S1-treated to -untreated fibers was determined. Data are displayed as means ± s.d. from two independent experiments.

**Supplementary Figure 7 (associated with Fig. 7). Induction of genomic uracil sensitizes UNG-dependent cancer cells to ATRi.**

(A) H1299 and H1838 cells were transfected with indicated siRNAs and treated with indicated concentrations of ATRi (VE-821) for 5-7 days. Cell viability was determined using CellTiter-Glo and normalized to the cells untreated with ATRi under each condition. Data are shown as mean ± s.d. (n=2 independent experiments).

(B) H1299, H1838, and HL-60 cells were treated with indicated concentrations of PMX and ATRi (AZD6738) for 5-7 days. Cell viability was determined using CellTiter-Glo and normalized to the cells untreated with ATRi under each condition. Data are shown as mean ± s.d. (n=2 independent experiments).

(C) Representative images and mean weight of excised tumors on day 36 following the treatment as in Fig. 7C. Scale bar, 1 cm.

(D) Changes in mouse weights during the treatment as in Fig. 7C. Body weights were measured every week. Data are shown as mean ± s.d.

(E) MGH 707.1, MGH 134.1, and MGH 143-3B cells were transfected with control or UNG#1 siRNA for 48 h. Levels of UNG2 in whole-cell extracts were analyzed by western blot. α-Tubulin is shown as the loading control.

(F) Cells as in (E) were analyzed by U-comet assay. Dot plots represent comet tail moment in individual cells, and bars display the mean in cell populations (n>50 cells per condition). One of two independent experiments is shown.

(G) Cells as in (E) were analyzed by DNA fiber assay. DNA fibers were treated with S1 nuclease (20 U/mL) for 30 min at 37°C, and the length of IdU replication tracts (n=125 fibers per condition) was measured. One of two independent experiments is shown.
